# “FDA-approved carbonic anhydrase inhibitors reduce Amyloid β pathology and improve cognition, by ameliorating cerebrovascular health and glial fitness”

**DOI:** 10.1101/2022.07.19.500681

**Authors:** Elisa Canepa, Rebecca Parodi-Rullan, Rafael Vazquez-Torres, Begona Gamallo-Lana, Roberto Guzman-Hernandez, Nicole L. Lemon, Federica Angiulli, Ludovic Debure, Marc A. Ilies, Leif Østergaard, Thomas Wisniewski, Eugenio Gutiérrez-Jiménez, Adam C. Mar, Silvia Fossati

## Abstract

Alzheimer’s disease (AD) is a devastating neurodegenerative disorder with no effective cure. Cerebrovascular and neurovascular pathology are early and causal hallmarks of AD, where cerebral amyloid angiopathy (CAA), the deposition of amyloid β (Aβ) at the cerebral vasculature, is present in about 90% of cases. Our previous work has uncovered the protective effect of carbonic anhydrase (CA) inhibition against Aβ-mediated mitochondrial dysfunction, production of reactive oxygen species (ROS) and apoptosis in vascular, glial and neuronal cells in culture. Here, we tested for the first time in a transgenic model of AD and cerebrovascular amyloidosis, the TgSwDI mice, a therapeutic regimen employing the FDA-approved CA inhibitors (CAIs), methazolamide (MTZ) and acetazolamide (ATZ). These drugs are used in humans for glaucoma, high altitude sickness, and other disorders, and can cross the blood-brain barrier. We found that both CAIs were non-toxic, significantly reduced cerebral amyloidosis, vascular, microglial and astrocytic Aβ accumulation, and ameliorated cognition. MTZ and ATZ treatment prevented caspase-3 activation in endothelial cells, microglia and astrocytes, reverted capillary constriction and microhemorrhages, reduced gliosis, and induced glial pro-clearance pathways, which are likely responsible for the reduction of Aβ deposition. Notably, we unveiled a critical new druggable target, revealing that the mitochondrial isozyme CA-VB is specifically upregulated in TgSwDI mouse brains, as well as in human brains of CAA and AD (with CAA) patients. Importantly, Aβ challenge induced CA-VB overexpression in human cerebral endothelial cells, and CA-VB silencing, mimicking CAIs effects, reduced Aβ-mediated endothelial apoptosis. This work paves the way for the application of CAIs in clinical trials for AD and CAA and uncovers CA-VB as a mediator of cerebral amyloid toxicity.

## Introduction

Alzheimer’s disease (AD) is a progressive neurodegenerative disorder without an effective cure. Morpho-functional alterations of the vasculature are critical pathological hallmarks of AD, that strongly contribute to the neurodegenerative process [1–7]. Importantly, cerebrovascular and neurovascular unit (NVU) changes occur at early stages, suggesting that cerebrovascular dysfunction (CVD) may contribute to the initiation and progression of the neuronal demise, and indicating a tight association between CVD, inflammation, neurodegeneration and cognitive impairment [1, 8–13]. Parenchymal amyloid beta (Aβ) plaques and tau neurofibrillary tangles (NFT) are considered the hallmark neuropathological features of AD. However, about 90% of AD cases also exhibit cerebral amyloid angiopathy (CAA), the deposition of Aβ aggregates within the vessel walls of arteries and surrounding the brain microvasculature [4, 14–21]. CAA can also be found in many non-AD older people, in a percentage correlating with age. Current immunotherapeutic approaches have also been shown to increase CAA and induce Aβ-related vascular abnormalities, such as ARIA (amyloid related imaging abnormalities) [22, 23]. Within the NVU, endothelial and glial cells actively co-operate in mediating vascular fitness and brain clearance, promoting the removal of toxic material from the brain, including Aβ. The modulation of systems implicated in Aβ degradation and perivascular clearance has recently aroused a lot of interest, especially in regard to their impact on the deposition of Aβ at the brain vasculature [4, 19, 24–26]. Indeed, if on one hand vascular Aβ deposition triggers CVD and affects brain clearance, on the other hand, CVD and dysregulated clearance exacerbate vascular Aβ deposition, leading to a vicious cycle, which boosts cellular stress, blood-brain barrier (BBB) permeability, metabolic waste accumulation, neuroinflammation, infiltration of blood-borne cells [27–29], and ultimately leads to neurodegenerative processes and cognitive dysfunction [1, 30, 31]. Yet, the molecular players mediating these harmful effects are not fully elucidated, and the scientific community urges to find novel targets and therapeutic strategies to prevent cerebrovascular and NVU impairment and to improve brain clearance in AD and CAA. We have previously demonstrated that Aβ induces mitochondrial dysfunction and caspase-mediated apoptosis in endothelial, glial and neuronal cells [15, 16, 32–35]. We also showed, through *in vitro* studies and in mouse brains injected with Aβ, that these detrimental mitochondrial stress and cell death phenomena are hampered by acetazolamide (ATZ) and methazolamide (MTZ), two pan-carbonic anhydrase inhibitors (CAIs), unveiling for the first time a possible contribution of carbonic anhydrases (CAs) to Aβ-initiated neurovascular cells impairment [36–38]. CAs catalyze the reversible hydration of CO_2_ to bicarbonate and a proton and are essential for pH regulation among many other cellular and physiological functions. Humans express 15 CA isoforms with different distribution patterns at organ/tissue and cellular level, including CA-VA and VB in the mitochondria. ATZ and MTZ CAIs are FDA-approved for non-AD-related conditions, such as glaucoma, high-altitude sickness, high-altitude cerebral edema and seizures; thus – advantageously – their pharmacokinetics and side effects are reported. It is known that the drugs are safe for long-term administration, can cross the BBB [37, 39–42], and that acute inhibition of CAs promotes cerebral blood flow (CBF), vasoreactivity and neuronal excitability in humans [43, 44]. Our recent studies demonstrated that ATZ and MTZ suppress mitochondrial dysfunction and apoptosis induced by Aβ in vascular and neural cells by reducing mitochondrial ROS production and loss of mitochondrial membrane potential [36, 38]. However, the effects of CAIs and the participation of CAs in Aβ-mediated cerebrovascular dysfunction, inflammation, Aβ deposition/clearance and neurodegeneration have never been dissected *in vivo* in a transgenic animal model of amyloidosis. In this study we tested for the first time in the AD/CAA field, a chronic therapeutic regimen employing the CAIs MTZ and ATZ in TgSwDI mice, which express human Amyloid-β Precursor Protein (hAPP), carrying the Swedish, Dutch and Iowa mutations. This model is characterized by extensive vascular Aβ deposition, starting at about 6 months of age, diffused parenchymal Aβ deposits and neuroinflammation. We assessed cognitive function and Aβ pathology following ATZ or MTZ treatment. Additionally, we tested the hypothesis that CAIs can halt or reduce Aβ neurovascular toxicity and promote vascular and glial cell fitness. Importantly, we assessed if specific mitochondrial CA isoforms mediate these protective neurovascular effects.

Overall, this work demonstrates for the first time that these CAIs can foster cerebrovascular fitness, revert the neuroinflammatory state, promote Aβ clearance and prevent cognitive impairment in a model of cerebral amyloidosis. Additionally, we provide evidence that points to a specific mitochondrial CA enzyme as a key player in CAA and AD.

## Results

### Chronic CAI treatment ameliorates spatial memory, reduces brain Aβ deposits and decreases caspase-3 activation

We have recently demonstrated for the first time in the AD field that CAIs reduce mitochondrial dysfunction and cell death pathways induced by Aβ in vascular and neural cells in culture, as well as in brain cells *in vivo* after intra-hippocampal Aβ injection [36–38]. Considering that mitochondrial health is essential for proper brain cell function, and that both mitochondrial impairment and CVD are very early events in AD pathology, we hypothesized that a treatment with CAIs may improve cognitive and pathological outcomes in a transgenic mouse model of cerebral amyloidosis with CAA, the TgSwDI mice. The FDA-approved CAIs ATZ or MTZ (20mg/kg/day) were chronically administered in the chow, which was otherwise identical to the control mouse diet (TestDiets). The treatment with ATZ or MTZ was started when Aβ deposition is mild in this model (7/8 months of age), to replicate a clinical treatment in MCI (Mild Cognitive Impairment) patients, or at a later age (12 months), to replicate clinical treatment in mild/moderate AD patients.

#### Behavioral studies

Once the mice reached a stage of advanced pathology (15/16 months), they all underwent a battery of behavioral tests. Before the cognitive assessment, weight was measured, and all study groups were subjected to motor activity tests, to verify that the CAI treatment was not toxic and did not affect motor skills. Rotarod performance (**Fig. 1A)** was impaired in all Tg groups compared to WT animals (TgSwDI p=0.016, Tg+ATZ p=0.012, Tg+MTZ p=0.006). Untreated TgSwDI mice had lower all-limb grip strength than WT control mice (p=0.003) (**Fig.1A**). However, grip strength in CAIs-treated Tg animals was not statistically different from WT animals. Forelimb (only) grip strength showed a similar pattern to the all-limb results (**Supplementary Fig.1A**). No significant differences in body weight due to the genotype or CAI treatment were observed between groups (**Fig. 1A**) or between same-gender groups (**Supplementary Fig.1A**), although as expected, males were heavier than females. Death rates also did not change between untreated Tg and CAI-treated Tg animals (**Supplementary Fig. 1A**), confirming that chronic CAI treatment was not toxic. Spatial memory was assessed using the Barnes maze test. First, all treated animals were compared to untreated Tg group and WT animals. The Barnes maze probe test data are presented in **Fig 1B**. Post-hoc comparisons between groups on each measure revealed that untreated TgSwDI mice ran a longer distance (p=0.0104) and made more mistakes (p=0.0117) than the WT control mice, indicating cognitive impairment. Tg mice treated with CAIs were not statistically different from WT animals, indicating that CAI treatment preserves spatial memory in TgSwDI mice (**Fig. 1B**). The same pattern of results was observed with ANOVA applied to the untransformed data or when using nonparametric Kruskal Wallis analysis. The results were similar after splitting the groups treated for 8 and 4 months (**Supplementary Fig. 1B**). To help rule out the potential contribution of confounding factors on Barnes maze performance measures, we also performed ANOVAs including body weight, distance travelled in the open field arena and percentage time spent in the center of the open field (not shown) as covariates. These covariates were not significant predictors of performance in any of the analyses (p > 0.05).

**Figure 1:**
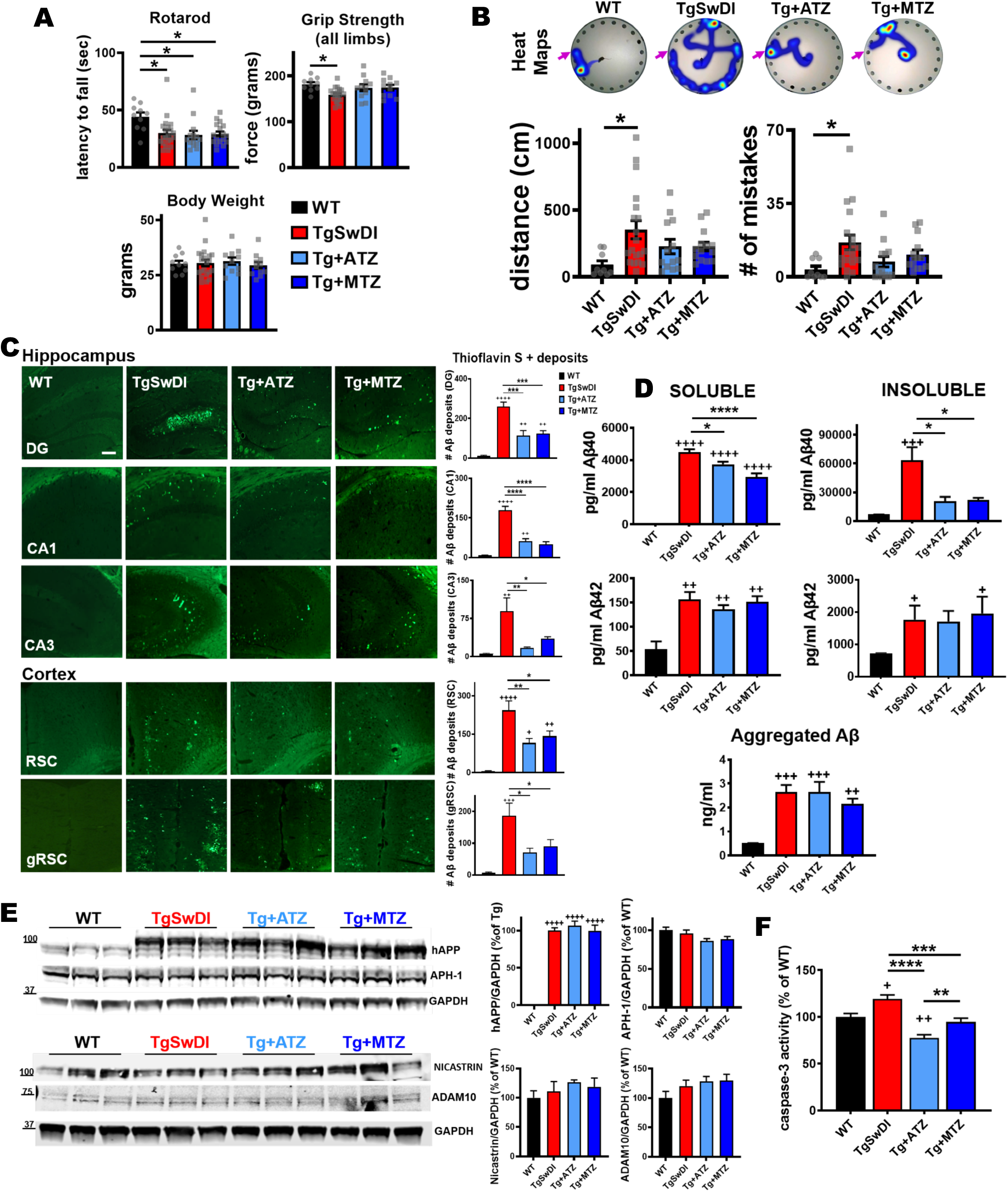
CAI treatment attenuates cognitive impairment, reduces brain Aβ pathology and decreases caspase-3 activation in TgSwDI mice. A) CAIs did not affect motor coordination tested with rotarod. In the rotarod performance, there was a significant main effect of group (F(1,57)=4.863, p=0.004) with body weight significant as covariate (F(1,57)=6.059, p=0.017). Post-hoc tests showed that all Tg groups were significantly impaired relative to WT control mice (TgSwDI p=0.016, Tg+ATZ p=0.012, Tg+MTZ p=0.006). The analysis of motor function tested with grip strength (all limbs) showed that there was a significant main effect of group (F(3,57)=5.449, p=0.002) on grip strength with body weight significant as covariate (F(1,57)=29.302, p=0.000). Post-hoc comparisons revealed that untreated TgSwDI mice had lower all-limb grip strength than WT mice (p=0.003). Body weight did not change between groups. B) Spatial memory tested via Barnes maze task in 15/16-month-old WT and TgSwDI mice, in the presence or absence of MTZ- or ATZ-treatment. The heat maps show the path (in blue) covered by the animals to reach the escape hole (indicated with pink arrows) in the probe test. The plots represent the distance covered (cm) and the number of mistakes made before finding the escape hole, during the probe test. There was a significant main effect of group on appropriately transformed indices of distance (F(3, 48)=4.456, p=0.008) and mistakes (F(3, 48)=5.059, p=0.004). Post-hoc comparisons between groups on each measure revealed that only untreated Tg mice were impaired in distance (p=0.0104) and mistakes (p=0.0117), compared to WT animals. Same pattern of results was observed with ANOVA applied to the untransformed data or when using nonparametric Kruskal Wallis analysis. WT: N=10, TgSwDI: N=19, ATZ: N=13 and MTZ: N=14. Two-way ANOVA and Dunn’s post-hoc test. Data are expressed as mean ± SEM. C) Representative images of cerebral Aβ deposits stained with Thioflavin S. 16-month-old untreated TgSwDI mice exhibit a greater amount of Aβ, in both hippocampus (DG, CA1 and CA3 areas) and cortex (retrosplenial and granular retrosplenial cortex, RSC and gRSC), compared to WT animals. CAI treatment significantly decreases Aβ deposits in all areas. Original magnification, 20x. Scale bar 150µm. On the right, relative quantification. WT, TgSwDI, ATZ and MTZ: N=5, n≥10 measurements acquired/group. D) Brain Aβ content measured by ELISA in soluble and insoluble fractions. Compared to age-matched WT, 16-month-old TgSwDI animals show significantly higher concentration (pg/ml) of both soluble and insoluble Aβ40 and Aβ42. CAI treatment diminishes soluble Aβ40, and even more considerably insoluble Aβ40, compared to Tg mice. Aggregated Aβ content (ng/ml) within the soluble fraction does not significantly change among Tg groups. Graphs represent N=3-8 animals/group. E) CAI treatment does not change the expression of hAPP and APP metabolism enzymes APH-1 and nicastrin (γ-secretase subunits), and ADAM10 (α-secretase). Quantification (right side), normalized vs GAPDH, and expressed in percentage vs WT mice (or % vs Tg in hAPP graph). N=3mice/group, n=9 technical replicates/group. F) Caspase-3 activity measured in brain homogenates. ATZ and MTZ treatments significantly reduce total cerebral caspase-3 activation, compared to untreated TgSwDI mice. N=3- 5 mice/group, n ≥12 measurements/group. In (C-F), one-way Anova and Tukey’s post-hoc test: * (vs Tg) and + (vs WT) p<0.05, ** and ++ p<0.01, *** and +++ p<0.001, **** and ++++ p<0.0001. Data are expressed as mean ± SEM.

Overall, these results indicate that spatial memory and grip strength are improved in CAI-treated Tg mice, without toxicity of the chronic treatment or effects on motor coordination.

#### Quantification of Aβ burden and APP processing

We next asked whether CAI-driven cognitive amelioration was associated with a reduction in cerebral Aβ burden. The mice were sacrificed after behavioral testing at 16 months, and brain sections were stained with Thioflavin S, which binds to fibrillar Aβ deposits. Fibrillar Aβ burden in hippocampal, cortical and hypothalamic regions was quantified using an unbiased sampling scheme and a semiautomated image analysis system. As expected, we found that 16-month-old TgSwDI animals presented extensive Thioflavin S+ Aβ deposits in the hippocampus (dentate gyrus [DG], CA1 and CA3 areas), and cortex (retrosplenial [RSC] and granular retrosplenial cortex [gRSC]) **(Fig. 1C**), as well as in the hypothalamus (**Supplementary Fig. 1C**). Aβ deposits were drastically reduced in all areas when Tg animals were treated with ATZ and MTZ from 8 to 16 months *(in DG, ATZ -56% and MTZ -52%; in CA1, ATZ -65.6 % and MTZ -72.6%; in CA3, ATZ -80.7% and MTZ -61%; in RSC, ATZ -52.4% and MTZ -41.6%; in gRSC, ATZ -62.5% and MTZ - 51.4%; in hypothalamus, ATZ -44% and MTZ -32%, vs Tg).* Differences in Aβ burden were also present in Tg mice treated for shorter time (from 12 to 16 months) with CAIs (**Supplementary Fig.1D**). However, longer treatment with ATZ and MTZ induced a much more evident reduction of cerebral Aβ pathology. Therefore, we focused our next analyses on the long-treatment group. We separated soluble and insoluble cerebral Aβ and measured the respective levels by ELISA (**Fig. 1D**). Aβ aggregates within the soluble fraction (corresponding to soluble oligomers) were also measured, using a specific aggregated Aβ ELISA assay. TgSwDI mice showed significant increase in all Aβ40 and Aβ42 species, compared to WT animals. In contrast to untreated Tg animals, CAI treatment significantly decreased Aβ40 content (the main peptide deposited in the vasculature, both in human AD and in this model), and remarkably reduced insoluble Aβ40 *(soluble Aβ40 reduction: ATZ -16% and MTZ -54.7%; insoluble Aβ40 reduction: ATZ -67.3% and MTZ -65%, vs Tg animals)*, suggesting that CAIs may reduce Aβ40 deposits around the cerebral vasculature (CAA). To rule out the possibility that this reduction was due to effects of CAIs on brain Aβ production through modulation of the expression of APP and/or its processing enzymes, we performed immunoblot analysis of these proteins. We found that CAIs did not alter the expression of human APP (hAPP) (**Fig. 1E**), or its amyloidogenic and non-amyloidogenic pathway enzymes, such as APH-1 and nicastrin (γ-secretase complex subunits) and ADAM10 (α-secretase), respectively (**Fig. 1E**), indicating that CAI-mediated reduction of cerebral Aβ deposition was not dependent on APP metabolism and Aβ production. This evidence, along with the knowledge that brain Aβ exists in a finely tuned balance between production and clearance, suggests that CAIs may promote Aβ clearance.

#### Total caspase-3 activation

To test whether the drastic cerebral Aβ reduction observed in CAI-treated mice was associated with a reduction in apoptotic cells, we measured the activity of caspase-3 in total brain lysates using a luminometric Caspase-Glo assay (**Fig. 1F**). We observed an increase in active caspase-3 in TgSwDI mice compared to WT, which was significantly decreased in ATZ- and MTZ-treated Tg animals, indicating that CAIs reduce the activation of apoptotic pathways in brain cells.

### ATZ and MTZ mitigate vascular Aβ burden, reduce endothelial active caspase-3, microhemorrhages, and revert vasoconstriction

To determine whether CAIs specifically reduce vascular Aβ deposition and Aβ-mediated endothelial apoptosis, we measured vascular Aβ and assessed caspase-3 activation in endothelial cells (ECs). First, immunofluorescence (IF) evaluation with an Aβ-specific antibody confirmed that ATZ- and MTZ-treated animals had massively decreased total cerebral Aβ burden, corroborating the Thioflavin staining. In addition, total caspase-3 activation, assessed by IF with an antibody that specifically binds the active form of caspase-3, was reduced in DG (**Fig. 2A**), cortex (**Fig. 2B**), and CA1 (**Supplementary Fig. 2A**), in CAI-treated animals *(in DG, ATZ -72% and MTZ -59%; in cortex, ATZ -58.5% and MTZ -59.7%; in CA1s, ATZ -65.7% and MTZ -57%,*

**Figure 2:**
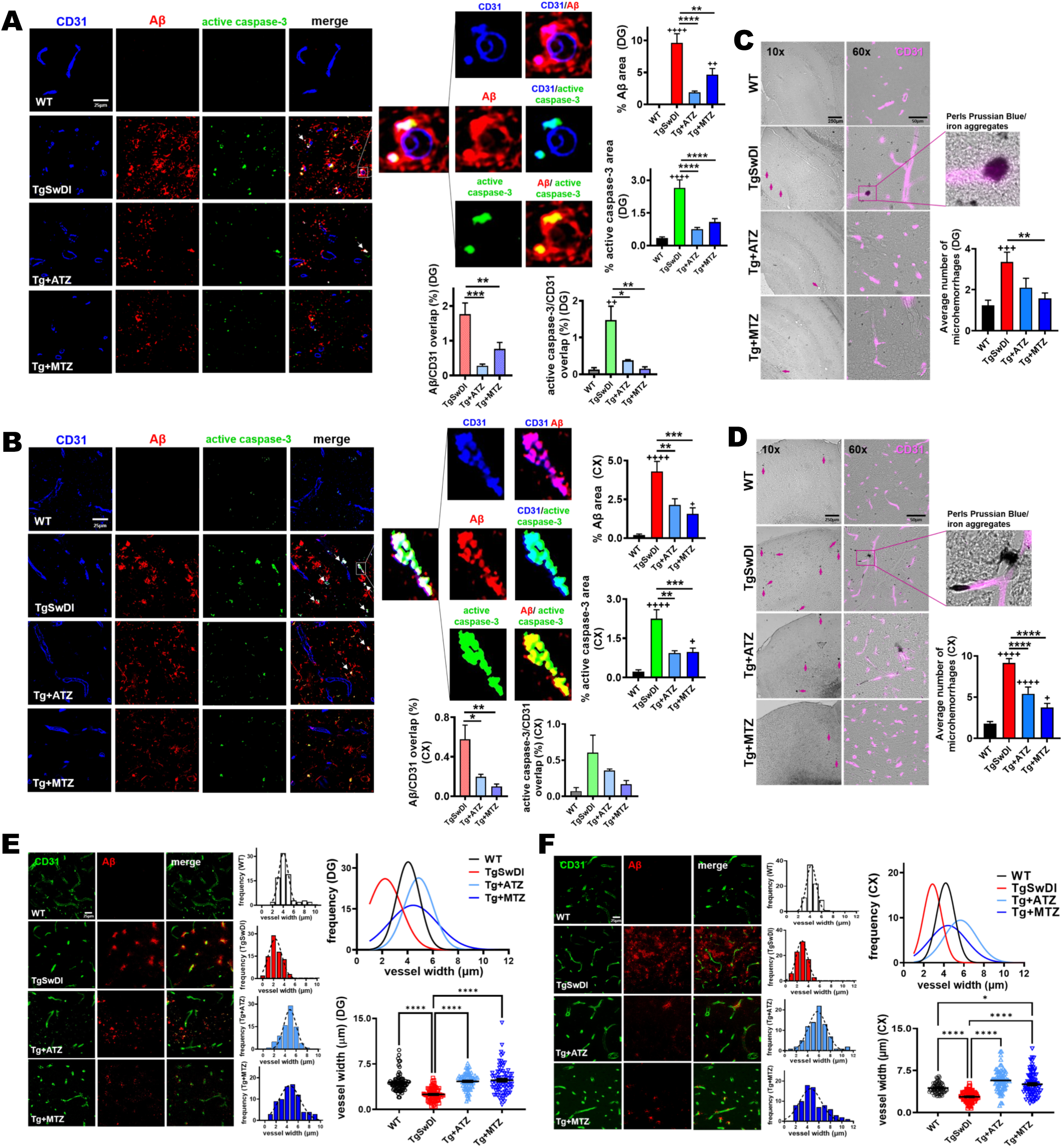
Vascular Aβ burden, endothelial caspase-3 activation, microhemorrhages and microvessel constriction are reduced by CAIs. A, B) Representative immunofluorescence images of DG (A) and cortex (B) of 16-month-old mice. TgSwDI mice present highly elevated cerebral Aβ (red) and active caspase-3 (green) staining, which are significantly decreased by CAI treatment. Original magnification, 60x. Scale bar, 25µm. On the right, relative quantification plotted as the percentage of area occupied by Aβ or active caspase-3 per acquisition field. For %Aβ graph, WT, TgSwDI and MTZ: N=5, ATZ: N=4, n≥12 measurements acquired /group. For % active caspase-3 graph, WT, TgSwDI and MTZ: N=5, ATZ: N=3, n≥9 measurements acquired/group. In the merged images, arrows point to Aβ deposits surrounding ECs (CD31+ ECs in blue). The magnified images illustrate overlap between the signals: ECs and Aβ (signal overlap in magenta), ECs and active caspase-3 (signal overlap in cyan), and Aβ overlapping active caspase-3 (yellow). Below, the graphs depict the percentage of Aβ and active caspase-3 signals, respectively, overlapping with CD31. For Aβ/CD31 overlap, N=5 for TgSwDI and MTZ, N=4 for ATZ, n≥12 measurements/group. For active caspase-3/CD31 overlap, N=3-5 mice/group, n ≥5 measurements/group. C) Representative 10x and 60x images of DAB-enhanced Perls Prussian Blue staining in DG of 16-month-old mice, showing higher number of microhemorrhages (MH) (indicated with arrows) in TgSwDI mice, vs WT animals, which are reduced in CAI-treated groups. Colocalization of iron aggregates with CD31+ BVs (magenta) is shown in the 60x and magnified images. Scale bars are 250µm and 50µm, respectively for 10x and 60x magnification. The plot on the right represents the average number of MH counted. WT, TgSwDI and MTZ: N=4, ATZ: N=3, n≥10 counts/group. D) Representative 10x and 60x images of Perls Prussian Blue staining, in cortex. MH presence (indicated with arrows) in TgSwDI mice, significantly decreased by CAI-diet. DAB-enhanced iron aggregates colocalizing with CD31+ BVs (magenta) shown in the magnified image. For magnification 10x and 60x, scale bar 250µm and 50µm, respectively. Relative quantification plotted on the right as the average number of MH. WT, TgSwDI and MTZ: N=4, ATZ: N=3, n≥10 counts/group. E) CAI treatment diminishes Aβ deposition (red) in DG, in TgSwDI mice, affecting vessel diameter (CD31, vascular marker, green). Original magnification, 60x. Scale bar, 25µm. On the right, vessel width frequency quantification in each group. On the top right, the frequency of all groups is plotted. On the bottom right, vessel diameter quantification, showing that CAI-fed mice, similarly to WT animals, displayed BVs with wider average diameter compared to untreated TgSwDI animals. WT, TgSwDI and MTZ: N=5, ATZ: N=4, n=80 vessels/group. F) CAI-diet increased blood vessel width in cortex, in 16-month-old TgSwDI mice. Original magnification 60x, scale bar 25µm. WT, TgSwDI and MTZ N=5, ATZ: N=4, n=80 vessels/group. In (A-F), one-way ANOVA and Tukey’s post-hoc test: * and + p<0.05, ** and ++ p<0.01, *** and +++ p<0.001, **** and ++++ p<0.0001. Data are expressed as mean ± SEM.

*vs Tg),* confirming the results of the luminometric caspase-3 activation assay. We then observed that vascular Aβ deposits (Aβ signal colocalized with CD31) also colocalized with active caspase- 3 signal in CD31+ ECs (**Fig. 2A** and **2B**; arrows and magnified images), suggesting that vascular Aβ triggers apoptotic processes in ECs, and supporting our previous *in vitro* studies [16, 32–35, 38]. Remarkably, ATZ and MTZ attenuated both microvascular Aβ burden (measured as Aβ staining overlapping with CD31 signal), and endothelial caspase-3 activation (active caspase-3 signal overlapping with CD31), in DG (*ATZ -74.2% and MTZ -89.4%, vs Tg;* **Fig. 2A**) and to a lesser degree (*ATZ -40.8%, p= 0.57, and MTZ -72.5%, p= 0.078, vs Tg*) in the cortex (**Fig. 2B**), indicating that CAIs mitigate both vascular Aβ deposition and Aβ-driven endothelial cell apoptosis. The presence of Aβ around the brain vasculature, especially in familial disorders or animal models bearing the Dutch or analogue vasculotropic mutations, is typically associated with cerebrovascular lesions, including microhemorrhages (MH) [45–48] . In TgSwDI mice, MH are present starting at 12 months of age [49]. Therefore, we tested whether the reduction of vascular Aβ and endothelial caspase-3 activation by CAIs could also result in a reduction of MH in the Tg mice. Compared to age-matched WT, 16-month-old Tg animals had a significantly higher number of MH in DG (**Fig. 2C**), cortex (**Fig. 2D**), and leptomeningeal arteries (**Supplementary Fig. 2B**). The number of MH was reduced in MTZ- and ATZ-treated groups *(in DG, ATZ -37.4%, and MTZ -53.2%; in cortex, ATZ -41% and MTZ -59%; in leptomeningeal arteries, ATZ -90.9% and MTZ - 93.7%, vs Tg),* indicating that CA inhibition prevents Aβ-mediated vessel wall degeneration. It has also been shown that Aβ deposition reduces vessel size in AD patients [50] and Tg mice [46, 51–53]. Hence, we measured vessel width in WT, Tg and treated animals. As expected, 16-month-old TgSwDI mice presented constricted vessels in DG (**Fig. 2E**) and cortex (**Fig. 2F**), compared to WT animals. In contrast, vessel diameter in CAI-treated Tg mice was not different from WT mice, corroborating our hypothesis that CAIs regulate cerebrovascular tone [54], besides promoting EC survival.

### Analysis of cerebral blood volume and flow in living animals

Previous studies have shown an age-dependent impairment in neurovascular coupling (NVC) in the TgSwDI model [55]. Therefore, we tested whether CAIs affected NVC impairment by measuring the relative changes in cerebral blood volume (rCBV) and cerebral blood flow (rCBF) in 10/11-month-old awake-restrained mice following whisker stimulation (**Fig. 3** and **Table 1**). Based on a time-series recording, we estimated the maximum evoked-response in CBV (rCBV Peak) and CBF (rCBF Peak), and extracted the area under the curve (rCBV A.U.C. and rCBF A.U.C.), which is considered as the total CBV and CBF change during functional activation/stimulation. Upon activation, we observed the expected increase in rCBV and rCBF in all groups (**Fig. 3B** and **3E**), but no significant difference in rCBV Peak (**Fig. 3C**), rCBV A.U.C. (**Fig. 3D**) and rCBF Peak **(Fig. 3F**), between Tg mice and WT animals was evident at this age. Interestingly, we observed a significant increase in rCBF A.U.C. in the MTZ-treated animals compared to the WT mice (Slope ≈ 0.42 ± 0.13, t(66)=3.16, p=0.002) (**Fig. 3G**).

Time series analysis of OISI (rCBV) and LDF (rCBF) M: Males, F: Females. **p > 0.01

**Table 1. Time series analysis of OISI (rCBV) and LDF (rCBF)**
| Variable | WT | Tg-Ctrl | Tg-ATZ | Tg-MTZ |
| --- | --- | --- | --- | --- |
| **Mice included** | 16 mice<br>(8 M and 8 F) | 17 mice<br>(6 M and 11 F) | 17 mice;<br>(11 M and 6 F) | 16 mice<br>(8 M and 8 F) |
| **OISI (N value)** | N = 30 | N = 32 | N = 27 | N = 30 |
| **rCBV Peak [%]** | 1.49 ± 0.49 | 1.56 ± 0.81 | 1.33 ± 0.62 | 1.57 ± 0.60 |
| **AUC [a.u.]** | 0.07 ± 0.04 | 0.09 ± 0.06 | 0.08 ± 0.04 | 0.1 ± 0.04 |
| **LDF (N value)** | N = 32 | N = 34 | N = 32 | N = 34 |
| **rCBF Peak [%]** | 10.87 ± 4.10 | 13.19 ± 5.44 | 12.27 ± 4.90 | 14.37 ± 6.27 |
| **AUC [a.u.]** | 0.50 ± 0.26 | 0.75 ± 0.42 | 0.70 ± 0.36 | 0.92 ± 0.45** |

**Figure 3:**
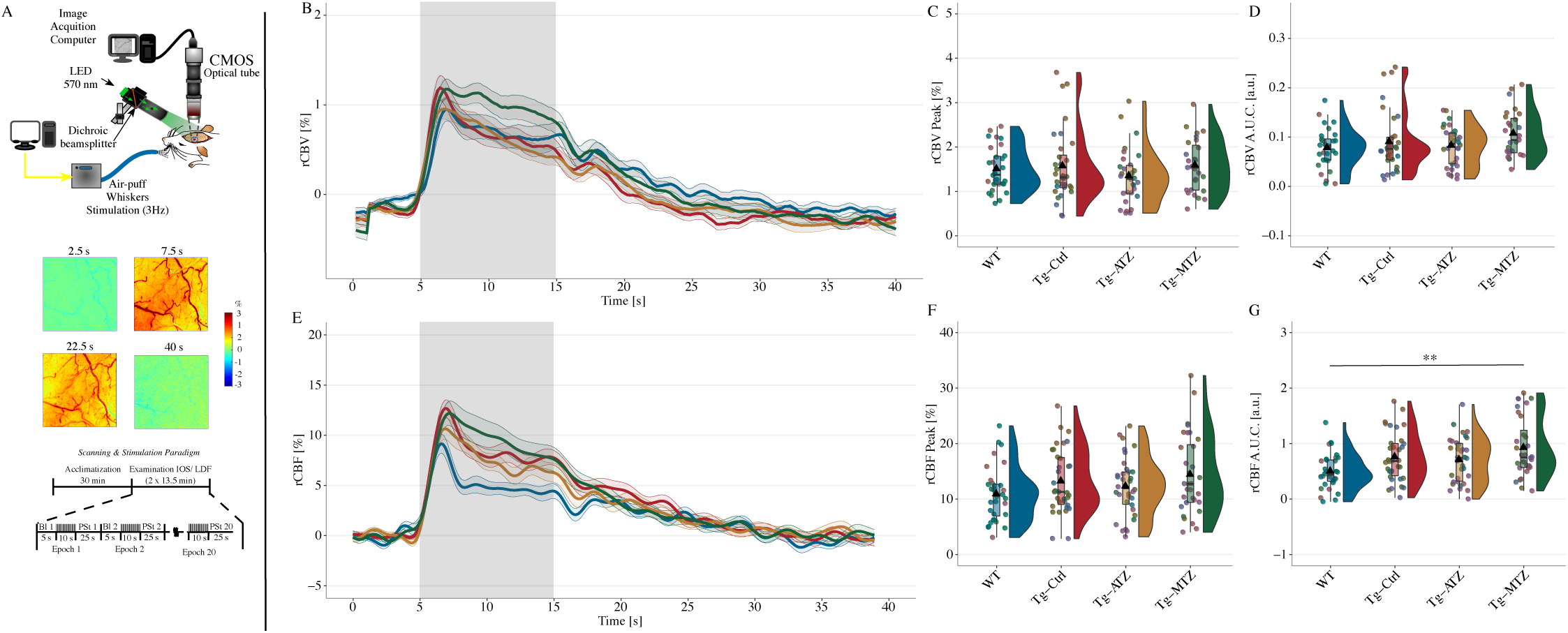
Analysis of cerebral blood flow and volume during functional activation. **A)** Experimental setup for IOS to measure total Hemoglobin (Hb), a proxy for cerebral blood volume (CBV). A LED constantly illuminated the cranial window with a spectrum of 570 ± 2nm (top), while AVI videos were recorded using a CMOS camera at 5-fps. IOS relies on the reflectance of light from the illuminated cortex. At 570nm, both oxy- and deoxy-Hb are absorbed at the same rate, equivalent to total Hb (rCBV). Whisker stimulation increases CBV, reducing the reflectance, hence the signal. In the postprocessing, the signal was inverted for better representation (middle). For IOS and LDF, whisker stimulation occurred from the second 5 to the second 15 in each epoch (bottom). **B)** Relative changes in CBV to the baseline (rCBV [%]) were estimated and plotted as time-series. **C)** No difference between the groups in the maximum CBV response (rCBV Peak [%]) to stimulation, and area under the curve (rCBV A.U.C. [a.u.]) (**D)**, during the stimulation period (10’). **E)** Estimated changes in regional cerebral blood flow (CBF) during functional activation, using a commercial LDF. **F)** No difference between groups in the maximum response to functional activation (rCBF Peak [%]). **G)** Analysis of the A.U.C. for regional CBF (rCBF A.U.C. [a.u.]). Tg+MTZ mice showed an overall increase in CBF response during the stimulation period, compared to the WT mice. N values: WT = 16 mice (8 males and 8 females); TgSwDI, N=17 (6 males and 11 females); Tg+ATZ, N=17 (11 males and 6 females); Tg+MTZ, N=16 (8 males and 8 females). ** p<0.01. Abbreviations: Bl: Baseline, PSt: poststimulation. Statistics: Cluster analysis using linear mixed-models predicting the changes in peak and A.U.C. by stimulation and adjusted for gender.

### CAI treatment reduces astrocytic Aβ accumulation and caspase-3 activation, rescuing Aβ- dependent astrogliosis

Astrocytes exert multiple functions in the brain, including mediating vascular fitness and permeability. They are physically in close association with both blood vessels (BVs) and neuronal cells [56], they assure the modulation of vascular response based on neuronal activity (NVC), and mediate perivascular clearance pathways [4, 26, 57, 58]. Moreover, both astrocytes and microglia act as scavenger cells, capable of internalizing and degrading brain waste molecules, including Aβ. Dysfunctional glial cells result in impaired Aβ clearance, and exacerbate cerebral Aβ overload and inflammatory state, also contributing to vascular Aβ deposition and loss of cerebrovascular functionality. In turn, vascular alterations, such as the observed Aβ-induced endothelial cell stress, can mediate glial activation and inflammatory state, and interfere with glial and perivascular clearance. Accordingly, we observed that severe Aβ accumulation in untreated Tg mice was associated with substantial astrogliosis in the hippocampus. Notably, both Aβ and Glial Fibrillary Acidic Protein (GFAP) staining were reduced by ATZ and MTZ treatment (**Fig. 4A**). Colocalization Aβ and GFAP signals showed that in Tg mice Aβ abundantly accumulated inside the astrocytes, while CAI treatment prevented the astrocytic Aβ overload (**Fig. 4B**). As expected, TgSwDI animals had an increased percentage of GFAP+ fluorescent area in the DG (**Fig. 4C**), indicating astrogliosis and hypertrophy of astrocytic processes. This pro-inflammatory phenotype was prevented by CAI treatment, in association with a drastic reduction of Aβ accumulation within astrocytes (quantified as colocalization of Aβ signal overlapping with GFAP signal). Concurrently, we observed a significant mitigation of active caspase-3 signal within GFAP+ astrocytes **(Fig. 4C**), particularly with ATZ treatment. ATZ and MTZ treatment resulted in a similar reduction of GFAP+ area, astrocytic Aβ accumulation and astrocytic caspase-3 activation in the cortex (**Fig. 4D**) and in the CA1 hippocampal area (**Supplementary Fig. 3A**). These results suggest that CAIs limit the activation of apoptotic pathways in astrocytes, facilitate Aβ removal and reduce astrogliosis. To corroborate these findings, we measured astrocytic hypertrophy, as the average area (µm^2^) of single GFAP+ cells in the hippocampus, cortex (**Fig. 4E**), and hypothalamus (**Supplementary Fig. 3B**). TgSwDI mice displayed a dramatic increment in the average single astrocyte area *(in DG, +260%; in CA1, +310%; in CA3, +322%; in RSC, +287%; in gRSC, +290%; in hypothalamus, +303%, vs WT),* denoting a typical astrocytic reactive phenotype, which was significantly reverted by CAI treatment *(in DG, ATZ -34% and MTZ -46.6%; in CA1, ATZ - 57% and MTZ -67%; in CA3, ATZ -39% and MTZ -37%; in RSC, ATZ -50% and MTZ -36.8%; in gRSC, ATZ -57% and MTZ -35%; in hypothalamus, ATZ -59.5% and MTZ -37%, vs Tg mice).* Overall, this evidence highlights the ability of CAIs to mitigate Aβ-initiated cerebral astrocytosis, and Aβ-driven apoptotic pathways in astrocytes, indicating that CAIs may improve glial health, favoring Aβ clearance.

**Figure 4:**
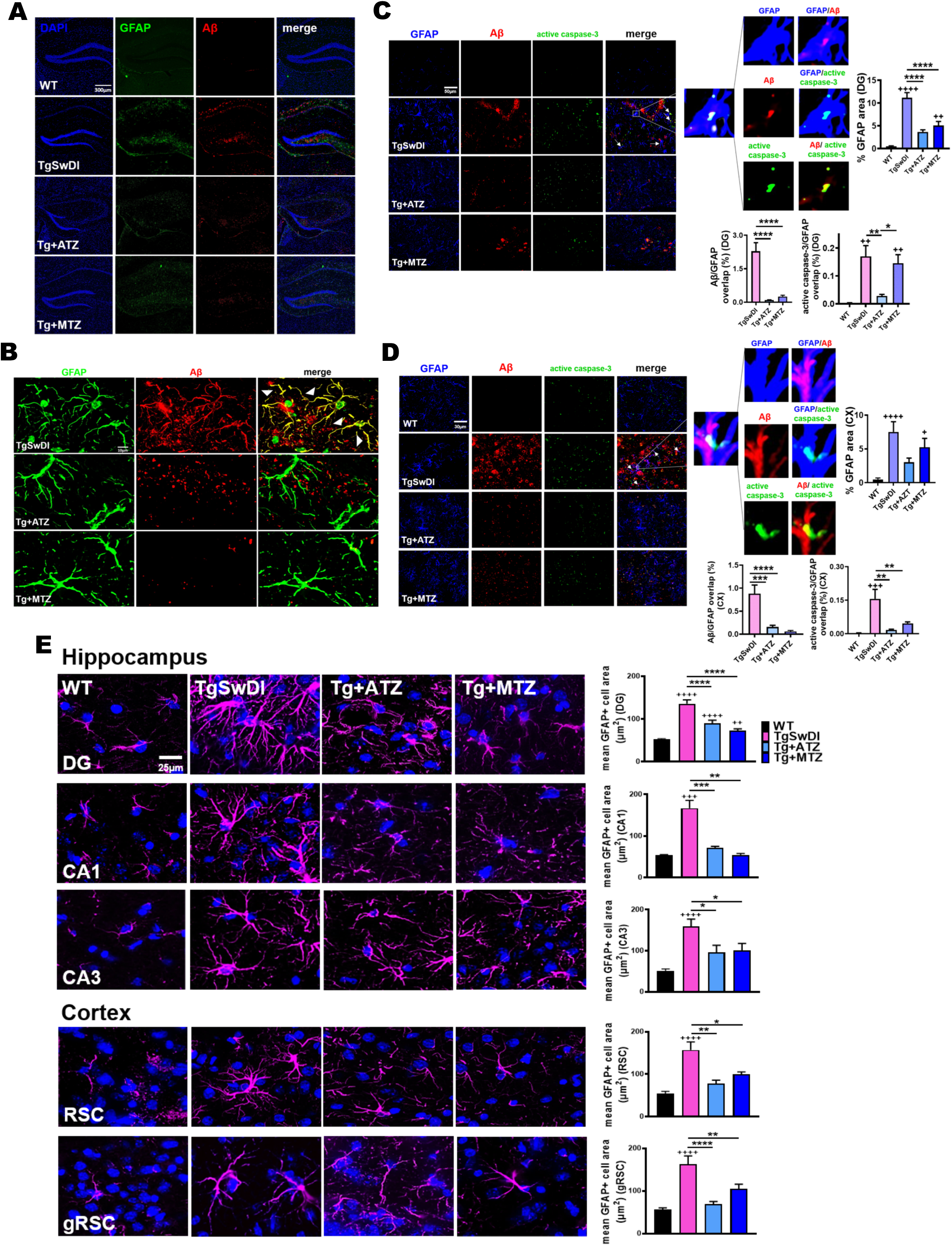
Astrocytosis, astrocytic Aβ content and caspase-3 activation are reduced in CAI- treated TgSwDI mice. **A)** 10x images of DG of 16-month-old TgSwDI mice show augmented GFAP expression (green) and Aβ overload (red). Nuclei stained with DAPI (blue). CAI treatment reduces both astrogliosis and Aβ deposits. Original magnification, 10x. Scale bar, 300µm. **B)** In TgSwDI mice, Aβ (red) is trapped within astrocytes (green), as shown by the colocalization (yellow) in the merged image. ATZ and MTZ treatment reduces astrocytic Aβ content. **C, D)** Representative immunofluorescence images of DG (**C**) and cortex (**D**). Compared to WT, untreated Tg animals exhibit greater expression of the astrocytic marker GFAP (blue), as well as Aβ (red) and active caspase-3 (green), all decreased by treatment with CAIs. Original magnification, 60x. Scale bar, 30µm. On the right, astrogliosis is plotted as %GFAP area per acquisition field. WT, TgSwDI and MTZ: N=5, ATZ: N=3, n≥9 measurements acquired/group. Arrows indicate colocalization of Aβ and active caspase-3 in astrocytes. The magnified images display Aβ within GFAP+ cells (signals overlap in magenta), astrocytic caspase-3 activation (signals overlap in cyan), and Aβ colocalizing with caspase-3 (yellow). Below, graphs represent the percentage of Aβ and active caspase-3 signals overlapping with GFAP+ cells, indicating that CAIs significantly reduced astrocytic Aβ accumulation and caspase-3 activation (significant for ATZ in DG, and for both CAIs in cortex) in astrocytes. WT, TgSwDI and MTZ: N=5, ATZ: N=3, n≥9 measurements/group. **E)** 60x magnified images representing astrogliosis in 16-month-old mice. TgSwDI brains are characterized by significantly increased GFAP+ average cell area (µm^2^) (GFAP in magenta), in both the hippocampus (DG, CA1 and CA3 areas) and cortex (RSC and gRSC). Nuclei are stained with DAPI (blue). 8-month-CAI-diet attenuates astrogliosis. Original magnification, 60x. Scale bar, 25µm. On the right, relative quantification of the mean area of one GFAP+ cell, for each brain area analyzed. For both hippocampus and cortex, WT, TgSwDI and MTZ: N=5, ATZ: N=4 animals/group. In **(A-E)**, one-way ANOVA and Tukey’s post-hoc test: * and + p< 0.05, ** and ++ p<0.01, *** and +++ p<0.001, **** and ++++ p<0.0001. Data are expressed as mean ± SEM.

### CAs inhibition attenuates microglial Aβ accumulation and caspase-3 activation, fostering a microglial pro-healing phenotype

Microglia are the primary brain resident innate immune cells and the most active form of immune defense in the CNS [59]. As astrocytes, microglia have an important role in brain clearance. In a physiological state, microglia migrate towards the site of interest, remove dead cell debris, and promote both extracellular and intracellular degradation of unwanted toxic substances, including Aβ. A prolonged insult prompts the microglial pro-inflammatory phenotype, resulting in morphological and biochemical changes, such as augmented expression of ionized calcium binding adaptor molecule 1 (IBA1) protein. Indeed, we found that 16-month-old TgSwDI mice had an increased percentage of IBA1+ signal area (**Fig. 5**), indicative of microgliosis, compared to age-matched WT animals *(in DG, +11fold change, F.C.; in cortex, +15 F.C.; in CA1, +8.3F.C., vs WT).* Importantly, CAI treatment significantly reduced IBA1 staining in DG (**Fig. 5A**), cortex (**Fig. 5B**), and in the CA1 hippocampal area (**Supplementary Fig. 4A**) (*in DG, ATZ -54% and MTZ -49%; in cortex, ATZ -64.7% and MTZ -56.7%; in CA1, ATZ -48% and MTZ -53%, vs Tg)*.

**Figure 5:**
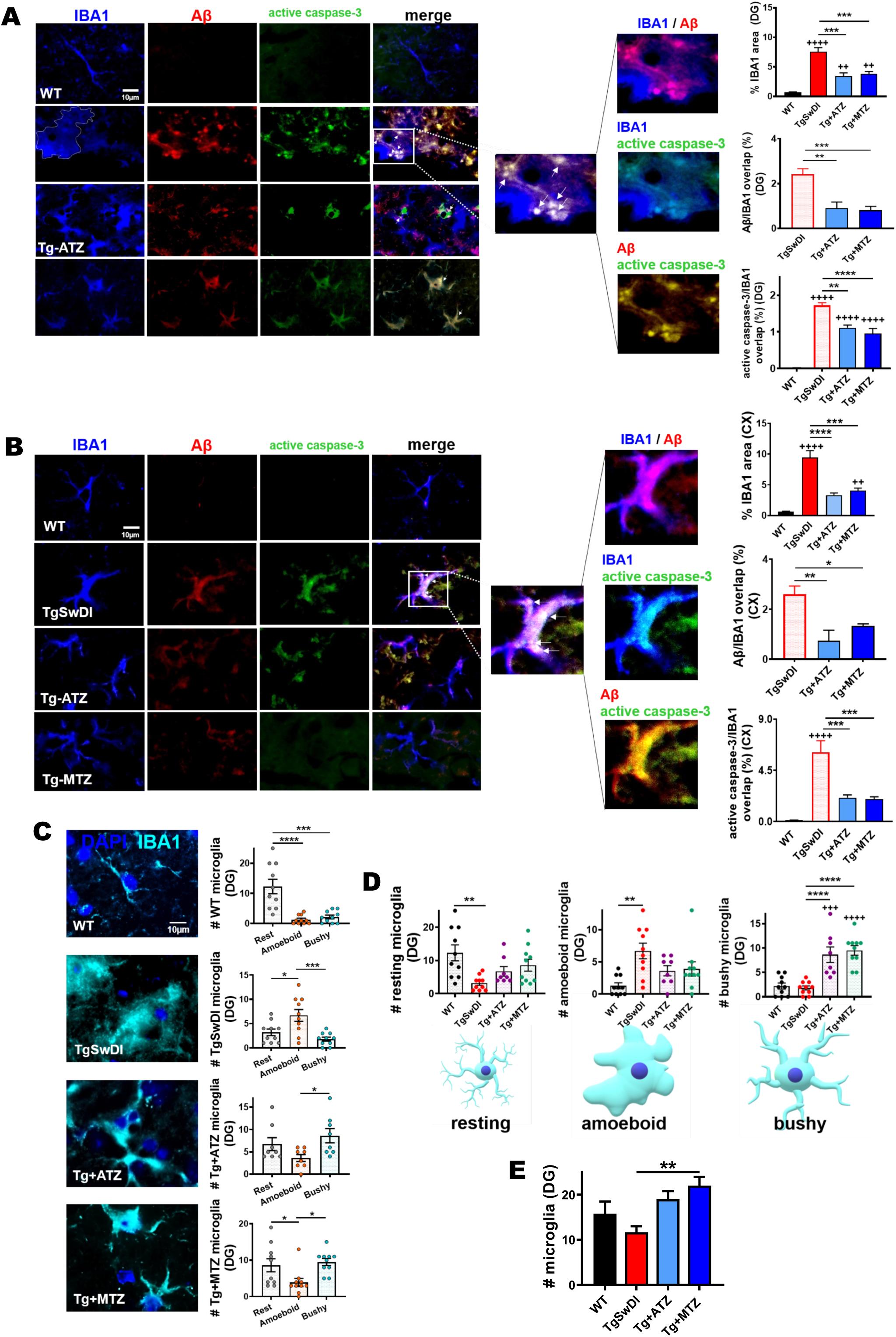
ATZ and MTZ reduce microglial Aβ overload and caspase-3 activity, and promote microglial pro-healing phenotype. **A, B)** Representative immunofluorescence images of DG (**A**) and cortex (**B**), showing that Tg animals exhibit increased microgliosis (IBA1, microglia marker in blue), Aβ (red) accumulation and active caspase-3 (green) in microglia, all rescued by CAIs. Original magnification, 60x. Scale bar, 10µm. On the right, % of IBA1 area per acquisition field. WT, TgSwDI and MTZ: N=5, ATZ: N=4, n≥8 measurements acquired/group. Arrows indicate microglia presenting internalized Aβ and active caspase-3. The magnified images show Aβ within IBA1+ cells (signal overlap in magenta), active caspase-3 within IBA1+ cells (signal overlap in cyan), and Aβ colocalizing with caspase-3 (yellow). On the right, plots represent the percentage of Aβ and active caspase-3 signals overlapping with IBA1+ cells, indicating that CAIs significantly reduce microglial Aβ overload and caspase-3 activation. WT, TgSwDI and MTZ: N= 5, ATZ: N=4, n≥8 measurements/group). **C)** Representative immunofluorescence images of microglia (IBA1 marker, cyan) for analysis of resting, amoeboid and bushy morphology. On the right, plots represent the different microglial phenotypes in each treatment group, in DG. WT mice have resting microglia as the most numerous subpopulation. TgSwDI have amoeboid microglia as most represented microglial type, while ATZ- and MTZ-treated mice present more bushy and resting microglia than amoeboid. WT, TgSwDI and MTZ: N= 5, ATZ: N=4, n≥8 measurements/group. **D)** Comparison of DG resting, amoeboid or bushy microglia between the different groups. The number of resting microglia is significantly higher in WT mice compared to Tg mice, but is not significantly different from WT in MTZ- and ATZ-treated mice. The amount of amoeboid microglia is the highest in Tg mice, while bushy microglia is more abundant in CAI-treated mice compared to WT and Tg. WT, TgSwDI and MTZ: N=5, ATZ: N=4, n≥8 measurements/group. **E)** Microglial cell (IBA1+ cells) count in DG. TgSwDI mice have fewer microglia than WT animals, while CAIs increase microglial number in Tg mice. WT, TgSwDI and MTZ: N=5, ATZ: N=4, n≥8 measurements/group. In **(A-E)**, one-way ANOVA and Tukey’s post-hoc test: * p< 0.05, ** and ++ p<0.01, *** and +++ p<0.001, **** and ++++ p<0.0001. Data are expressed as mean ± SEM.

In order to elucidate whether this change was associated with Aβ overload and active caspase-3 in microglia, we analyzed the amount of Aβ and active caspase-3 signals overlapping with IBA1 staining. We found that untreated Tg mice displayed a massive microglial Aβ accumulation in DG (**Fig. 5A**), cortex (**Fig. 5B**), and CA1 (**Supplementary Fig. 4A**), accompanied by a robust microglial-specific activation of caspase-3 (**Fig. 5A**, **5B** and **Supplementary Fig. 4A**) *(in DG, +63F.C.; in cortex, +49.5F.C.; in CA1, +100F.C., vs WT)*. Notably, CAIs attenuated both Aβ deposition and caspase 3 activation in microglia (**Fig. 5A**, **5B** and **Supplementary Fig. 4A**) *(for Aβ, in DG, ATZ -62% and MTZ -66%; in cortex, ATZ -71% and MTZ -48%; in CA1, ATZ -62.9% and MTZ -86%, vs Tg. For caspase-3 activation, in DG, ATZ -35% and MTZ -44.5%; in cortex, ATZ -65% and MTZ -67.6%; in CA1, ATZ -69% and MTZ -89%, vs Tg)*. We further investigated CAI impact on neuroinflammatory responses by evaluating microglial shape, which has been accepted as an indicator of their functional state [60]. Resting microglial cells present ramified morphology with some long and fine processes [61]. In the event of an insult, microglia become active and rapidly change their morphology from ramified to bushy cells with multiple short, thickened and sturdy processes [62]. At the site of injury, a pro-healing microglial phenotype contributes to neuroprotection [63–65]. Under excessive and chronic stress, microglia keep modifying their shape, retracting their processes, and adopting an amoeboid shape [66], which is indicative of a severe pro-inflammatory state associated to the release of neurotoxic factors. In each treatment group, resting, bushy and amoeboid microglia subpopulations were present (**Fig. 5C**). As expected, resting microglia were the most represented subpopulation in the WT group (**Fig. 5C**). In contrast, in untreated TgSwDI brains, amoeboid microglia were the most abundant type. Notably, ATZ- and MTZ-treated mice were predominantly characterized by bushy microglia (**Fig. 5C**). These data suggest that CAIs reduce the toxic pro-inflammatory phenotype and promote the active but neuroprotective phenotype. When plotting resting, amoeboid and bushy microglia in the different treatment groups in DG (**Fig. 5D**), resting microglia were significant higher in WT compared to TgSwDI mice, and partially rescued by ATZ and MTZ. As expected, amoeboid microglia were higher in Tg mice, and partially reduced by MTZ and ATZ. Interestingly, the number of bushy microglia was significantly higher in CAI-treated animals, compared to both WT and untreated Tg mice, suggesting that CAIs may contribute to promote a microglial pro-clearance active phenotype. In the cortex, we found that TgSwDI animals had a significantly higher number of bushy microglia, compared to amoeboid, with similar levels in CAI-fed animals **(Supplementary Fig. 4B**), likely indicating a higher resilience of microglia in the cortical areas of these Tg mice. Both ATZ- and MTZ-treated groups presented a significantly higher number of resting microglia in the cortex compared to TgSwDI mice (**Supplementary Fig. 4C**), with similar counts as in WT animals, but the treatment did not affect the total number of amoeboid and bushy cells. To evaluate CAI effects on the total number of microglia, we counted IBA1+ cells. We found that TgSwDI mice had a trend to a reduced number of microglia in DG, compared to WT mice (although not significant), possibly due to microglial apoptosis, while CAIs rescued this loss, exhibiting microglial counts even higher than WT animals **(Fig. 5E**), suggesting a prevalence of heathy activated microglia in CAI-treated Tg mice. In the cortex, all Tg mice showed a higher number of microglial cells, compared to WT (**Supplementary Fig. 4D**), in line with the microglial phenotypes described above. These results indicate that CAIs reduce Aβ-mediated microglial toxicity and microglial caspase activation, promote microglial pro-healing phenotypes, and reduce the neurotoxic inflammatory phenotype, particularly in the hippocampus.

### CAIs promote microglial and perivascular macrophage phagocytic activity

The triggering receptor expressed on myeloid-cells-2 (TREM2) is expressed by microglia and promotes microglial survival, proliferation and phagocytic activity [67–69]. In our TgSwDI mice, TREM2 levels appeared slightly lower *(-19% in DG and -16% in cortex, not significant)* compared to age-matched WT animals. However, ATZ and MTZ treatment significantly boosted TREM2 expression compared to untreated Tg mice in the DG, with a similar trend in the cortex (**Fig. 6A**). When compared to WT mice, ATZ also significantly enhanced TREM2 expression also in the CA1 hippocampal area (**Supplementary Fig. 5A**). The increase in TREM2 induced by CAIs suggests that these drugs promote microglial phagocytic activity and therefore Aβ clearance, in line with the observed reduction of total, vascular and glial Aβ deposits. To corroborate this hypothesis, we analyzed the presence of CD68, a microglial/perivascular macrophage (PVM) activation marker, located in the endosomal/lysosomal compartment, which plays a major role in the clearance of brain waste material [70]. IHC analysis revealed that TgSwDI mice exhibited increased CD68 levels in the DG (**Fig. 6B**, more than +2.5 F.C.), and even greater levels in the CA1 (+3.8 F.C.) (**Supplementary Fig. 5B**), in contrast to WT animals. Strikingly, in both areas, CAI treatment further increased CD68 expression compared to untreated Tg mice (in DG, ATZ- and MTZ-treated mice showed +1.65 and +1.44F.C., respectively, compared to Tg; in CA1, ATZ and MTZ groups had +1.7 and +1.26F.C., respectively, in contrast to untreated Tg). These data, in combination with the increase in TREM2, reinforce the hypothesis that CAIs may reduce Aβ deposition by promoting microglial/PVM phagocytic activity. We observed the presence of an elevated number of CD68+ cells that colocalize with vascular Aβ deposits in the perivascular spaces (**Fig. 6B**, DG, magnified images), suggesting that CAIs (particularly ATZ) may prompt PVM phagocytic activity, likely contributing to perivascular Aβ clearance. Indeed, the CAI-treated groups presented a reduced Aβ content within CD68+ cells (DG, **Fig. 6B**) compared to untreated Tg animals. These results may suggest an increased amount of active perivascular CD68+ cells migrating over the pathological deposits, accompanied by an increased degradation of Aβ, likely resulting in the observed reduction of vascular Aβ (**Fig. 2A**, **2B**, and **Supplementary Fig. 2A**). In the cortex (**Fig. 6C**), MTZ treatment particularly boosted the total CD68 signal compared to Tg mice. The percentage of Aβ signal within CD68+ cells was slightly reduced in CAI-treated mice, although it did not reach statistical significance. Similarly to what observed in the DG, ATZ induced a higher CD68 signal over Aβ deposits (**Fig. 6C**), suggesting an increased trophism of CD68+ cells towards Aβ degradation. Immunoblot analysis of CD68 expression corroborated the IHC assessment, showing increased CD68 levels in TgSwDI animals compared to WT, with an even greater expression in CAI-diet fed mice (**Fig. 6D**). Overall, these results suggest that CA inhibition promotes Aβ removal by microglia/PVM, facilitating Aβ clearance.

**Figure 6:**
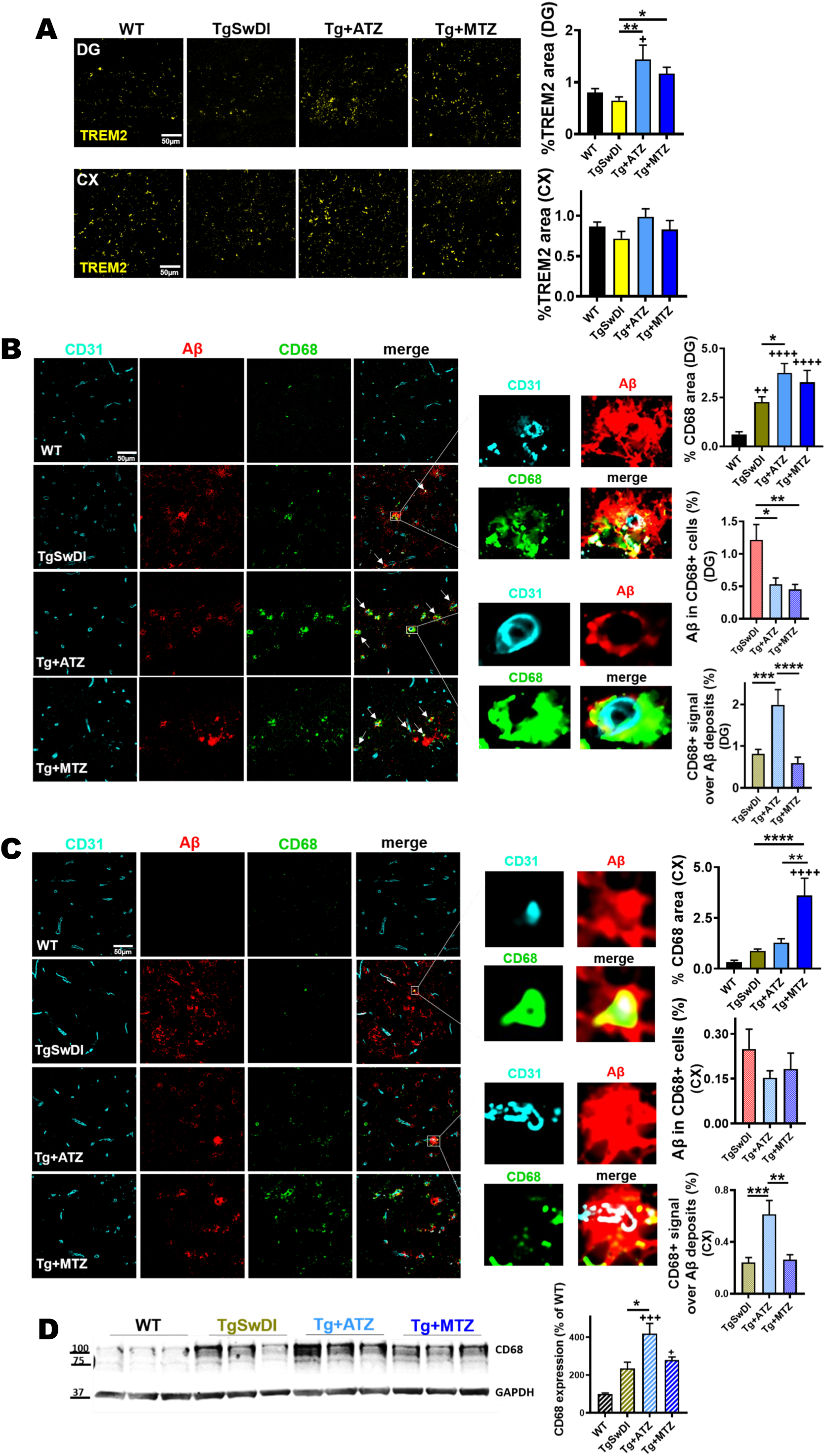
**CAI treatment increases TREM2 and CD68+ perivascular phagocytic cells. A**) ATZ and MTZ significantly increase microglial TREM2 expression in DG in TgSwDI mice, and show a trend to increase in the cortex (cx). Relative plots on the right. For % TREM2 area, both in DG and cortex, WT, TgSwDI and MTZ: N=5, ATZ: N=3, n≥9 measurements acquired /group. * and + p<0.05 and **p<0.01, One-way ANOVA and Tukey’s post-hoc test. **B, C)** Expression of CD68 (green) around microvasculature (CD31, cyan), and co-localization with Aβ (red) in DG (**B**) and cortex (**C**). Original magnification, 60x. Scale bar, 50µm. CAI-fed animals present higher phagocytic activity marker (CD68), plotted as %CD68 area per acquisition field, compared to WT and Tg mice. TgSwDI: N=7, WT: N=6, ATZ: N=4, MTZ: N=5, n≥12 measurements acquired/group. * p<0.05, ++ p<0.01, ++++ p<0.0001, One-way ANOVA and Tukey’s post-hoc test. Arrows indicate vascular Aβ internalized by CD68+ perivascular macrophages (PVM), as shown in yellow in the magnified images. On the right, colocalization plots for both Aβ within CD68+ cells (Aβ/CD68) and CD68+ area over Aβ deposits (CD68/Aβ). TgSwDI: N=7, MTZ: N=5, ATZ: N=4, n≥11 measurements/group. For Aβ/CD68 colocalization, * p<0.05 and ** p<0.01. For CD68/Aβ colocalization, *** p<0.001 and **** p<0.0001, ++++P<0.0001. One-way ANOVA and Tukey’s post-hoc test. **D)** WB of total brain homogenates showing that CD68 expression is significantly increased by CAIs, particularly ATZ. N=3/group, * and + p<0.05, +++ p<0.001, One-way ANOVA and Tukey’s post-hoc test. Data are expressed as mean ± SEM.

### CA-VB is a mediator of Aβ-driven neurovascular dysfunction

We have shown above that CAI treatment inhibits caspase-3 activation in ECs, astrocytes and microglia in Tg mice. We have previously demonstrated in cellular models that CAIs reduce caspase activation in cells composing the NVU by preventing Aβ-driven mitochondrial dysfunction and mitochondrial ROS production (particularly H_2_O_2_) [16, 32–36, 38]. Among the 15 known human CA isoforms, CA-VA and -VB are expressed solely in the mitochondria, and CA- II (a widespread cytosolic isoform abundant in the CNS) translocates to the mitochondria during aging and neurodegeneration [71]. Because mitochondrial mechanisms are responsible for the protective effects exerted by CAIs, we tested whether changes in mitochondrial CA isoforms were present in TgSwDI mice. Immunoblot analysis unveiled that, compared to WT, 16-month-old TgSwDI brains displayed a significant increased CA-VB expression (+1.75 F.O.C, p<0.01) (**Fig. 7A**), and, strikingly, CAI treatment restored CA-VB levels similar to WT brains (**Fig. 7A**). When we measured the cerebral levels of CA-VA and CA-II in our model, we did not observe significant alterations among the different groups (**Fig. 7A**), suggesting that specifically CA-VB overexpression may be pathologically induced by cerebral amyloidosis. To corroborate this result, we evaluated mitochondrial CA expression in CAA human cortices, and detected upregulated human CA-VB levels, with a strong trend to significance (127% increase, p=0.06; **Fig. 7B**), compared to the controls, and no changes in CA-VA and CA-II expression (**Fig. 7B**). Similarly, we found that CA-VB was overexpressed approaching significance (511% increase, p=0.06; **Fig. 7C**) also in human AD brains presenting CAA pathology (AD+CAA). Combining the human CAA and AD+CAA groups, we found a significant upregulation of CA-VB levels (**Fig. 7D;** 215% increase, p<0.05), suggesting that brain amyloidosis and CAA may trigger a specific CA-VB overexpression. To confirm this, and due to the importance of vascular dysfunction in our model, we evaluated the expression of CA-VB in human cerebral microvascular ECs challenged with 25µM Aβ40-Q22 (the vasculotropic Dutch mutant present in the TgSwDI mice) or 10µM Aβ42. In line with the expression pattern observed in the Tg animals and in the human subjects, we observed that Aβ, particularly Aβ40-Q22, induced an increase in CA-VB expression (**Fig.7E**), but not in that of CA-VA or CA-II (**Supplementary Fig. 6A and 6B**). Hence, to test the hypothesis that inhibition of endothelial CA-VB results in protective effects, we assessed microvascular EC apoptosis following Aβ challenge in the presence or absence of CA-VB silencing RNA (**Fig. 7F** and **7G**), and compared its effects to CA-VA and CA-II silencing (**Supplementary Fig. 6C** and **6D**). Silencing efficiency was confirmed via PCR after 48hrs (-80% CA-VB expression in siCA- VB conditions, compared to a scrambled siRNA control) (**Fig. 7F**). As expected, ECs treated with Aβ peptides at concentrations known to cause apoptosis in our models (10µM Aβ42 or 25µM Aβ40-Q22, for 24hrs after silencing) [16, 32, 38], underwent apoptosis in presence of the scrambled siRNA control, while CA-VB silencing significantly rescued Aβ42- and Aβ40-Q22- induced EC apoptosis (**Fig. 7G**). In contrast, when CA-VA and CA-II were silenced, we did not observe any attenuation of Aβ-induced apoptosis (**Supplementary Fig. 6C** and **6D**), pointing to CA-VB as a specific target for mitigating Aβ-induced EC stress, and thus the resultant neurovascular dysregulation, in AD and CAA. On the contrary, CA-VA downregulation, per se, induced apoptosis (**Supplementary Fig. 6C**), as expected, since CA-VA is considered an essential enzyme [72, 73]. Interestingly, CA-VA was also decreased in ECs treated with Aβ40-Q22 (**Supplementary Fig. 6A**). Overall, this set of experiments suggests that the protective properties of CAIs observed in our cellular and animal models may be mediated, at least in part, through the mitigation of a maladaptive Aβ-induced overexpression of the mitochondrial CA-VB isoform. Importantly, upon CAI treatment, both Aβ deposition and CA-VB overexpression were reduced in Tg mice. This finding, together with the fact that CA-VB silencing reduces Aβ-induced EC apoptosis, indicates that Aβ-induced CA-VB overexpression may be a mediator of cerebrovascular stress and cell death.

**Figure 7:**
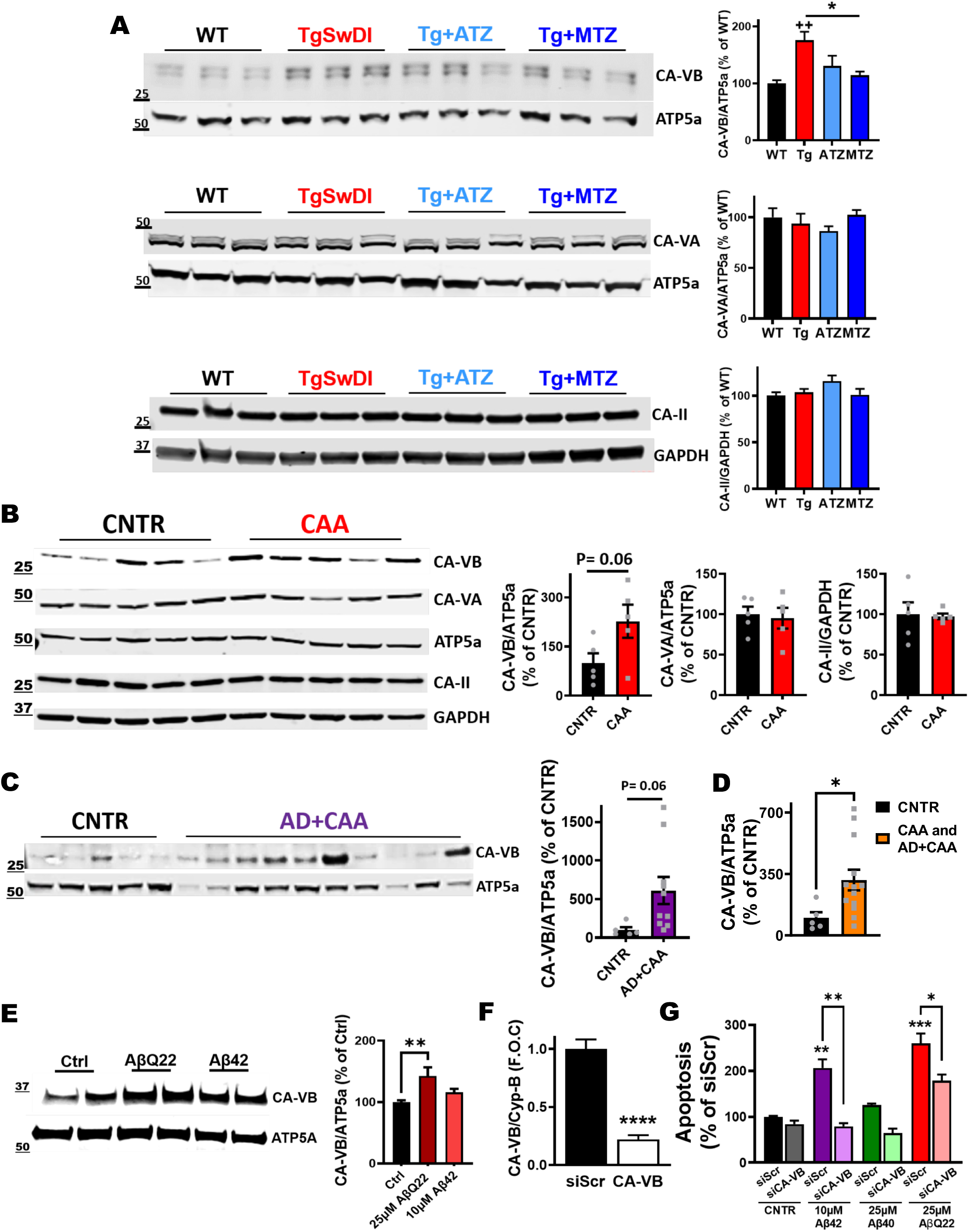
CA-VB expression increases in Tg mice, AβQ22-treated ECs and CA-VB silencing prevents endothelial apoptosis. **A)** Immunoblot for CA-VA, -VB and -II in total brain lysates of 16-month-old mice. TgSwDI brains display a significant increase in the mitochondrial carbonic anhydrase-VB (CA-VB) compared to age-matched WT mice, while CAI-treated brains do not show the CA-VB increase (CA-VB normalized to the mitochondrial protein ATP5a). The expression of CA-VA (normalized to ATP5a) and of CA-II (normalized to GAPDH) does not change. N=3 animals/group, n=6 technical replicates/group. * p<0.05 and ++ p<0.01, One-way ANOVA and Tukey’s post-hoc test. **B, C)** Biochemical analysis of mitochondrial carbonic anhydrases in human cortices. CA-VB expression is upregulated in CAA (+127%, p=0.06) **(B)** and in AD+CAA subjects (+511%, p=0.06) **(C)**, compared to healthy controls (CNTR). No significant alterations in CA-VA and CA-II levels, between CAA and CNTR brains **(B)**. CNTR and CAA groups N=5, and AD+CAA group N=10. Two-tailed unpaired t test. **D)** The combination of CAA and AD+CAA groups shows significant upregulation of CA-VB levels compared to CNTR. CNTR N=5; CAA group combined with AD+CAA group N=15. *p<0.05. Two-tailed unpaired t test. **E)** Western Blot analysis of CA-VB in cerebral ECs after 24hr challenge with Aβ40-Q22 (25µM) and Aβ42 (10µM). CA-VB was normalized to the mitochondrial protein ATP5a. Quantification is represented on the right. The expression of CA-VB is significantly increased following 24hrs Aβ40-Q22 treatment. Data represents the combination of at least three experiments each with 2 replicates, graphed as mean + SEM. One-way ANOVA and Dunnett’s post-test. **p< 0.01 **F)** qRT-PCR for mRNA expression levels of CA-VB and Cyp-B (housekeeping control gene) in cerebral ECs 48hrs post-transfection with siRNA for CA-VB (siCA-VB) or with a scrambled siRNA sequence (siScr) as control. **** p<0.0001 vs. siScr, Unpaired two-tailed t-test. **G)** CA-VB silencing prevents apoptosis, measured as the formation of fragmented nucleosomes (by Cell death ELISA^plus^), after challenge with Aβ42 (10µM), Aβ40 (25µM) or Aβ40-Q22 (25µM) for 24hrs (starting after the 48hr silencing). The graph displays one representative experiment of at least N=3 experiments, each performed in duplicate (n=2). * p<0.05 and ** p<0.01 vs. siScr control, One-way ANOVA, and Tukey’s post-hoc test. Data are expressed as mean ± SEM.

## Discussion

This study investigated, for the first time in the AD field, whether the FDA-approved CAIs ATZ and MTZ ameliorate amyloidosis, cognitive impairment, Aβ-initiated CVD, NVU pathology and neuroinflammation *in vivo*, in a transgenic model of cerebral amyloidosis with CAA, the TgSwDI mice. We demonstrated that a chronic treatment with both CAIs results in a reduction of Aβ pathology in all brain areas analyzed, decreasing in particular fibrillar Aβ40 species, the main component of CAA vascular deposits, with associated spatial memory and grip strength improvement. While preventing cognitive impairment, the two CAIs showed no toxicity, bearing no effects on body weight or death rates. TgSwDI animals treated with ATZ and MTZ showed reduced vascular, microglial and astrocytic Aβ accumulation and lower caspase-3 activation in endothelial and glial cells, together with an improvement of microvascular abnormalities and glial reactivity phenotypes.

The presence of neurovascular alterations at early stages of AD pathology points to CVD as one of the major causal players in the disease onset and progression. Many clinical studies, including the Alzheimer’s Disease Neuroimaging Initiative (ADNI), suggest the involvement of cerebrovascular disease very early in AD, either in the development or as a result of Aβ deposition [7, 74]. Recent transcriptomic studies also revealed selective vulnerability of vascular cell subpopulations associated with clinically diagnosed AD [75].

The two investigated CAIs can cross the BBB, are safe for chronic systemic administration in humans, and are used clinically for prolonged periods against glaucoma, high-altitude sickness, related high-altitude cerebral edema and other non-AD-related indications, including seizures [37, 39, 40, 42]. We were the first to show that ATZ and MTZ reduce Aβ-induced loss of mitochondrial membrane potential, mitochondrial H_2_O_2_ overproduction, caspase-9 and -3 activation and Aβ- mediated apoptosis in cerebral vascular, neuronal and glial cells in culture and in the hippocampus of WT mice injected with Aβ [36–38].

In this study, when applying the drugs for the first time in a transgenic model, we observed a clear reduction in Aβ deposition in the hippocampus, cortex and hypothalamus in CAIs-treated Tg mice compared to age-matched untreated Tg mice, which does not appear to be due to a dysregulated APP processing. The behavioral assessment revealed that 15-16-month-old TgSwDI mice had lower grip strength and rotarod performance relative to age-matched WT controls, suggesting impairments in limb strength, motor learning or coordination. TgSwDI also showed impaired performance in the probe test of the Barnes maze (increased distance and errors), independent of observed changes in motor function, implicating deficits in learning and memory. Chronic treatment with both CAIs shifted performance on the majority of these tests. The transgenic mice treated with ATZ and MTZ were not significantly different from WT in grip strength (all limbs and forelimbs) and Barnes maze distance and errors. These findings suggest that CAIs can act to ameliorate peak limb force and spatial memory function. Neither ATZ nor MTZ exerted detectable changes on rotarod performance, which suggests specificity in the functional circuits impacted. CAIs may thus exert little to no impact on striatal and cerebellar function, but could act to improve the cortical and/or hippocampal deficits associated with impairments in motor strength and mnemonic function that have been reported in TgSwDI mice, as well as other AD mouse models, and are consistent with our findings of a reduced Aβ pathology.

It is widely accepted that accumulation of Aβ in the brain parenchyma and around cerebral vessels occurs when its production and clearance are unbalanced. Extracellular Aβ can be removed from the brain by multiple clearance systems [4, 26], such as the intramural periarterial drainage (IPAD) pathway [4, 76, 77], along brain artery and capillary basement membranes, due to the contractility of vascular smooth muscle cells [25, 78], or the astroglial-mediated CSF-ISF bulk flow, known as glymphatic clearance [79–81]. The meningeal lymphatic system provides a complementary clearance route for cerebral waste products, which recently gained interest in neurodegenerative disorders, such as AD [82, 83]. Glial cells such as astrocytes and microglia are essential for removing Aβ through phagocytosis and degradation, which, when efficient, prevent Aβ accumulation [84–87] . Astrocytes are also essential to support vascular function. Their end-feet wrap the blood vessels, aiding endothelial junction protein stabilization and CBF regulation [88]. They provide physical and metabolic support to the brain, regulating brain clearance [89] and maintaining the brain antioxidant system [90, 91]. Reactive astrogliosis is an early process in AD and CAA pathogenesis [92–94], and multiple studies suggest that astrocytic Aβ clearance is impaired in AD settings [84, 95, 96]. Microglia, upon prolonged insults, prompt to pro-inflammatory phenotype, resulting in morphological and biochemical changes, which perpetuate brain inflammatory responses [97, 98], while hindering clearance and degradation of unwanted products. All these clearance processes may act in parallel or in synergy to preserve brain homeostasis [24], and all appear to be dependent on vascular fitness and functionality, glial cell health, or both [4, 26, 99]. In his study, we show that the treatment with CAIs improves both cerebrovascular and glial fitness.

We observed that in this model, cerebral amyloidosis is associated with caspase activation in ECs, microglia and astrocytes. Caspase activation, whether or not resulting in apoptotic cell death, is known to decrease the ability of vascular and glial cells to perform their metabolic functions, such as the above-mentioned clearance mechanisms [100, 101]. Caspase activation has also been linked to neurotoxic signaling to neurons, and to tangle formation [101–104]. Here, CAI treatment robustly decreases caspase activation in ECs, astrocytes and microglia in Tg mice, likely improving vascular and glial cells ability to perform their physiological functions.

Importantly, we tested whether CAIs improve cerebrovascular health and functionality. We showed that CAIs reduce the number of cerebral microhemorrhages, which characterize human Dutch and Iowa CAA cases [45–48], often occur in late onset AD and CAA brains [105–107], and are present starting at 12 months in this animal model [49]. Additionally, while untreated TgSwDI mice showed significant vasoconstriction in DG and cortex, vessel diameter in CAI-treated mice was restored to WT levels. These results corroborate the impact of CAA on EC degeneration and blood vessel dysfunction, as observed in AD subjects [50]. Furthermore, they confirm that CAIs may be able to regulate cerebral vessel tone, possibly by both reducing EC mitochondrial stress and ROS production, as we have previously shown [36–38, 42], as well as through their known potential to stimulate vasodilation [54, 108, 109]. It is known that in acute ATZ vasodilatory tests, 1000mg ATZ achieves a supramaximal dilative effect for 20’ [108]. It is important to note that the doses we used chronically in our Tg model, when translated to human doses (182mg/day for a 70kg person after allometric scaling), are many folds lower than the doses used in the acute ATZ challenge. Interestingly, when we assessed CBF and CBV response during functional activation in TgSwDI mice we did not see significant effects in the maximum CBF or CBV response (or in their respective AUCs) compared to WT animals. Similarly, Tg mice fed with CAIs at our doses for 6 months did not differ significantly from untreated Tg animals, indicating that chronic effects of low doses of CAIs on CBF and CBV may be different from acute effects of high doses of the drugs, typically used for the ATZ vasodilatory challenge.

It is well accepted that increased oxidative stress mediated by Aβ in ECs and neural/glial cells promotes mitochondrial dysfunction, leading to caspase activation and eventually apoptosis [16, 110–112]. We have shown that CAIs prevent mitochondrial dysfunction, mitochondrial ROS production, caspase activation and cell death in multiple brain cell types, including endothelial and glial cells [35, 36, 38, 113, 114]. Therefore, we postulate that a reduction of EC and glial mitochondrial stress and caspase activation is the primary mediator of the observed effects, leading to the reduced Aβ pathology and improved cognition, likely by ameliorating vascular and glia-mediated clearance pathways. Accordingly, we show that CAIs decrease both astrocytic and microglial Aβ content, together with reduced caspase-3 activation and reduced microglial and astrocytic reactivity markers, suggesting that glial cells in the treated animals are healthier and can better degrade Aβ. Indeed, healthy glial cells are essential in the brain. It was shown that microglial ablation in AD triggers vascular hemorrhage and exacerbates Aβ deposition in brain vasculature [115]. Moreover, microglia react to inflammation augmenting vascular interactions [116], thus regulating BBB integrity following an inflammatory insult [117]. Reactive microglia also induce capillary permeability by producing ROS, and promoting the migration of blood-borne inflammatory cells through the BBB [118, 119]. Thus, it is clear that preventing microglial and astrocytic stress and over-reactivity can also promote vascular health, and vice-versa [99], as we observe in our CAI-treated mice.

Interestingly, the reduction of Aβ and caspase-3 activation observed in microglia of CAI-treated mice was associated with reduced IBA-1 expression and amoeboid shape, but with an augmented number of total microglia. Human studies show that AD subjects present microglia with larger cell bodies and decreased process length and/or branching. In contrast, Aβ-immunized AD subjects have twice the number of microglia found in both AD and control groups, together with an increased number of ramified microglia and total process length [120], in line with the increased cell numbers and higher amounts of ramified or “bushy” microglia that we show in our CAI-treated animals. Interestingly, treatment with CAIs, especially ATZ, increases the levels of TREM2 in Tg mice. It is known that TREM2 promotes microglial phagocytosis and Aβ clearance by fostering the survival of activated microglia [69, 121, 122], and its variants are risk factors for late-onset AD [123]. TREM2 deficiency also reduces immunotherapeutic Aβ clearance [124], and loss of TREM2 is associated with early stages of CAA [94]. Therefore, the observed increase in TREM2 confirms that CAIs can facilitate microglial phagocytosis and, ultimately, Aβ clearance.

In addition, our work shows that CAI-treated Tg mice express high levels of CD68+ cells compared to untreated Tg animals and WT animals. PVM and microglia express the phagocytic marker CD68 [125, 126], which localizes to late endosomes and lysosomes [127], and is upregulated in response to inflammatory stimuli. CD68 preferential location within late endosomes suggests a role in peptide transport/antigen processing [128]. PVM are associated with Aβ deposits in meningeal and cortical BVs and contribute to vascular health [12, 129, 130]. Notably, ATZ-treated mice show increased CD68+ cells over Aβ vascular deposits, indicating that CAIs may promote Aβ degradation by PVM, resulting in reduced vascular Aβ. In line with our results, in TgCRND8 mice, PVM depletion by clodronate augmented the accumulation of vascular Aβ [130], and in J20 mice the ablation of the scavenger receptor class B type-I (SR-BI), expressed in PVM, triggered vascular and parenchymal Aβ deposition in the hippocampus, and exacerbated cognitive impairment [129]. Moreover, CD68 expressing cells surrounding 6E10+Aβ plaques were augmented when old AD and young WT slices were co-cultured, in association with a reduced Aβ plaque size [131]. In line with this data, the increase in phagocytic CD68+ PVM that we observed upon CAI treatment may improve the clearance of perivascular Aβ.

It needs to be noted that ATZ, at high doses, is also known to decrease CSF production [132], and appears to reduce glymphatic clearance in animal models [133]. However, we believe this is not a detrimental factor for our therapeutic strategy, as supported by our observed global reduction of cerebral Aβ deposition, and improvement in cognition, without signs of toxicity, after long-term administration of the drugs at the lower doses we propose. In confirmation of this, impaired glymphatic clearance is also present in normal pressure hydrocephalus (NPH) [134], and treatment with low-dose ATZ (125-375mg/day) in this disorder was shown to reverse periventricular white matter hyperintensities [135], suggesting that the protective effects of low doses of ATZ may be uncoupled from its effects on glymphatic clearance in this disorder, as well as in our model.

Furthermore, arterial pulsation, which is essential for multiple clearance mechanisms including IPAD [25, 136], can also be improved by CAIs [42, 54], suggesting that ATZ and MTZ could exert additional positive effects by improving vascular smooth muscle cell tone and function, which is currently the subject of further studies by our group.

Humans have 15 CA isoforms, differentially expressed at the tissue and cellular level [37, 42, 137]. Because mitochondrial mechanisms are responsible for the protective effects exerted by CAIs in our studies and others [16, 32–36, 38, 42, 138–140], we asked if the two mitochondrial CAs, CA- VA and CA-VB [138, 141], and CA-II, the most abundant CNS CA enzyme, which was found to localize at the mitochondria during aging and neurodegeneration [71], are affected in our model. Interestingly, we found that the mitochondrial CA-VB is specifically overexpressed in TgSwDI brains compared to WT and that CAI treatment reverts this effect. Most importantly, we found that CA-VB expression was upregulated also in human subjects affected by CAA or by AD with CAA, corroborating that cerebral Aβ-deposits may induce CA-VB overexpression, which may be a mediator of mitochondrial dysfunction and apoptosis [139, 140]. In line with this hypothesis, we demonstrated that CA-VB is specifically increased in ECs after challenge with vasculotropic Aβ. Strikingly, silencing of CA-VB, but not CA-VA or CA-II, prevents Aβ-induced EC apoptosis. This discovery points for the first time to CA-VB as a mediator of Aβ-induced cerebrovascular pathology, which may be, at least in part, responsible for the protective effects of CAIs. This finding also paves the way for future studies to understand the specific impact of CA-VB deregulation and its specific genetic or pharmacological modulation in CAA and AD, which are currently being developed in our laboratory.

Overall, our results underscore the substantial potential therapeutic impact of CAIs in AD and CAA, providing a mechanistic understanding for the protective effects of the inhibitors and paving the way to clinical trials to repurpose these FDA-approved compounds in MCI or early AD/CAA patients. Limitations of this study include the fact that this mouse model lacks tau pathology, another important hallmark of AD. Indeed, studies of the effects of CAIs in mouse models containing both Aβ plaques and hyperphosphorylated tau are currently in progress in our laboratory. Imaging (MRI and two-photon microscopy) studies are also in progress to extend our understanding of the effects of these drugs on the cerebral microcirculation, oxygen diffusion, brain structures/volumes, and perivascular clearance pathways. These studies, together with future clinical trials, will allow us to establish whether CAI treatment can be an effective pharmacological strategy to apply in AD/MCI patients and, potentially, in other AD-related dementias.

## Materials and Methods

### Animals and treatment

TgSwDI (APP-Swedish, Dutch, Iowa) mice (C57BL6/6 background) were obtained from Dr. Thomas Wisniewski (New York University, NYU), and bred internally. These animals carry the human APP gene (isoform 770) with the Swedish (K670N/M671L), Dutch (E693Q) and Iowa (D694N) mutations, under the control of the mouse neuronal Thy1 promoter, triggering an enhanced abnormal cerebral Aβ production and deposition. In particular, mutations within Aβ peptide sequence, such as the Dutch and the Iowa (at position 22 and 23, respectively), are mainly associated with fibrillar Aβ burden in the brain microvasculature and diffused parenchymal deposits. The animals were housed in accordance with Institutional and National Institutes of Health (NIH) guidelines, and the animal protocol was approved by the Animal Studies Committee. Mice were maintained under controlled conditions (∼22°C, and in an inverted 12hr light/dark cycle, lights OFF 10am-ON 10pm) with unrestricted access to food and water. To determine the impact of CAIs on Aβ-mediated pathology in TgSwDI mice, we fed animals for 8 months (8-16 months) or for 4 months (12-16 months) with a CAI-supplemented diet. MTZ or ATZ (20mg/kg/day, corresponding to 100 ppm), were incorporated in a control rodent diet (5053 by TestDiets, Quakertown, PA). To monitor for potential toxicity, the weight of each animal was measured every 2 weeks for the first 2 months, and before behavioral analysis, and skin/fur appearance was observed weekly.

### Behavioral analysis

Mice were tested in a behavioral test battery performed during the dark phase of the light cycle. The animals were transported and acclimated to the testing room, at least 30’ prior to each testing session. All behavioral testing was conducted in accordance with the NYU School of Medicine’s Institutional Animal Care and Use Committee, NYU School of Medicine.

#### Rotarod

Locomotor function, coordination, balance and motor learning were measured using an accelerating rotarod procedure across 3 trials spaced by a 40’ ITI. 5 mice were run simultaneously on 9.5cm diameter rods of a 5-lane rotarod apparatus (IITC Life Science Inc.). All mice were first given a habituation trial in which they were placed on static horizontal rods, and required to stay there, without falling, for 1’. Animals were then tested in 3 experimental trials, during which the rod was rotating and accelerating steadily from 4 to 40rpm over the course a 5’ period, and the latency to fall (sec) from the rods was recorded, and plotted as shown.

#### Grip strength

Paw/limb grip strength was assessed as the maximal horizontal force (gr) generated by the subject while grasping a specialized 6x10cm stainless steel grid platform connected to a sensitive force sensor (Bioseb). Two different grip strength indices were collected: forelimbs and all limbs (combined fore- and hind-limbs), and both were adjusted to the body weight. For both indices, each mouse was subjected to 6 testing trials with an inter-trial interval (ITI) of 10-20”, and a 40’ interval between forelimb and all-limb measurements. In each trial, the mouse was put onto the grid platform, allowing only its forepaws (forelimbs) or all four paws (all limbs) to clasp onto the central top-half portion of the grid. Once the paws were grasping the grid and the mouse’s torso was in horizontal position, the animal was moved to the cage by the operator. The truncated mean (highest and lowest scores removed) of 6 consecutive trials was taken as the index of grip strength. The body weight was measured after grip strength testing.

#### Barnes Maze

The Barnes maze procedure provides an index of visuospatial learning and memory in mice to navigate and escape an aversive, open area. The maze apparatus consists of a beige, textured plastic platform surface, 36” in diameter, elevated 91.5cm from the floor (San Diego Instruments, San Diego, CA, USA). The platform has 20 holes (5cm dimeter), equally spaced around the periphery of the platform (∼2.5cm from the edge). A gray plastic escape box (∼10x8.5x4cm) was located under one of the holes (target/escape hole), and kept consistent across trials. The spatial location of the target hole was counter-balanced across subjects/groups. Visible distal cues were placed around the room and remained constant throughout the testing period. The maze was illuminated by an overhead lamp (∼600 lux) above the center of the platform. Mice were tested for 12 trials across 6 consecutive days: one habituation trial (day 1), 10 training trials (days 1-5, 2 trials/day), and one probe trial (day 6). For the habituation trial, each mouse was placed into the center of the maze under an inverted, clear 500ml beaker for 1’. A white noise generator (∼80-90dB at platform level) was turned ON and, after 1’ period, the beaker was slowly moved to the target hole with the animal still inside. The mouse was allowed to enter the escape box and explore it for 2’. After the habituation (2hr ITI), each mouse was given 2 standard training trials each day, during which the animal was placed into the center of the maze in a plastic 15×15×20cm start box. The white noise was turned ON and, after 10/15”, the start box was lifted. The mouse was given a maximum of 3’ to find the designated hole and escape box. Once the animal entered the escape box, the white noise was turned OFF, and the animal left inside the box for 30’, and then moved to the home cage. In the case of failure of finding the escape hole within the 3’ limit, the mouse was slowly guided to the target hole under the inverted beaker, as described above. Each day, trials were spaced by 1.5/2hr ITI and, for each trial, the maze platform was rotated. On day 6, a 2’ probe trial was conducted, similarly to the training trials, except that the escape box was removed. Behavior was recorded using an overhead camera for later tracking, and the analysis performed by Noldus Ethovision software (v11.5). Distance travelled, number of mistakes (non-target holes visited) and latency to find the target hole were measured as indices of spatial memory, and then plotted as shown.

### Mouse brain processing

After behavioral analysis, brains were harvested for biochemical and immunohistochemical (IHC) assessments. Briefly, animals were anesthetized with pentobarbital, transcardially perfused with either solely ice-cold saline solution (0.9% NaCl) to wash out blood from vasculature, or with saline solution first, and then with 4% paraformaldehyde (PFA) to fix tissues, based on the experimental purpose [36]. Unfixed brains were removed, flash frozen in liquid N_2_, and stored at -80°C until their processing for the biochemical assessment. Fixed brains were left O/N at 4°C in 4% PFA, and incubated at 4°C one day in 15% sucrose solution, followed by additional 2 days at 4°C in 30% sucrose, as cryoprotectant. They were then washed in PBS, assembled in a plastic mold with Tissue-Tek O.C.T. compound (Fisher Scientific), and frozen with a mixture of liquid N_2_ and isopentanol. Brains were stored at -80°C until sectioning. Serial cryostat sections of 8µm thickness were collected on positively charged microscope slides (Fisher Scientific), and stored at −80°C until further immunohistochemical analysis.

### Thioflavin S staining

Brain slices were washed in dH_2_O and incubated in 0.015µM Thioflavin S for 30’, following which, sections were washed in 80% Et-OH. After washing in 1xPBS, slices were dried, and dipped into water-based medium before mounting [142]. All chemicals were from Sigma (St. Louis, MO). Thioflavin S staining was quantified in at least 4 evenly spaced coronal sections/animal and 5 animals/group using the ImageJ fluorescence analysis tool. Cortical (RSC and gRSC), hippocampal (dentate gyrus, CA1 and CA3) and hypothalamic Aβ fibrillar burden (defined as the number of Thioflavin+ deposits) was quantified separately.

### Extraction and quantification of soluble and insoluble Aβ

Mouse brains were homogenized in ice-cold homogenization buffer containing 20mM Tris pH 7.4, 1mM EDTA, 1mM EGTA, and 250mM sucrose. After homogenization, equal amount of protein was used to obtain the Aβ soluble and insoluble fraction. To obtain the soluble fraction, which contains Aβ monomers, oligomers, and protofibrils, a diethylamine (DEA) extraction was performed [143, 144]. Briefly, 0.4 % DEA buffer (v/v) was added to each sample and centrifuged at 50,000xg for 1hr at 4°C. The supernatant was transferred to a new tube, and 0.5M Tris-HCl was added at a 1:10 dilution. The resulting soluble fraction was then stored at -80°C for further analyses. To obtain the insoluble fraction, which represents the fibrillar Aβ associated to plaques/CAA, a formic acid (FA) extraction was performed. The resulting pellet was homogenized in 99% FA buffer. The sample was then centrifuged at 50,000xg for 1hr at 4°C, followed by the addition of the neutralization buffer containing 1M Tris base, 0.5M Na_2_HPO_4_, and 0.05% NaN_3_, and then stored at -80°C for further analyses. The soluble and insoluble fractions were used to measure the levels of human Aβ40 (Invitrogen) and Aβ42 (Invitrogen) by solid-phase sandwich ELISA according to manufacturer’s instructions. Aggregated Aβ within the soluble fraction was measured using a conformation-specific ELISA assay (Invitrogen).

### Caspase-3 activity measurement

Brains were homogenized in 25mM HEPES, 5mM MgCl_2_, 1mM EGTA buffer, containing 1X Halt protease inhibitors (Thermo Fisher), following the 100mg tissue/1ml homogenization buffer proportion. Cerebral homogenates were centrifuged at 13.000rpm for 15’ at 4°C, and supernatants were collected and stored at -80°C until usage. Following protein content quantification via BCA method, 10µg protein/sample were added to each well of a white-walled 96-well luminometer plate, in combination with Caspase-Glo® 3/7 Reagent (Promega) [33]. After incubating for 1hr at RT, the caspase-3 activation-dependent luminescence of each sample was read in a plate-reading luminometer, and plotted as percentage of caspase-3 activity of WT animals.

### Immunohistochemical assessment

For immunostaining evaluation, brain sections were blocked with 10% NGS, 1% BSA solution for 2hrs at RT, and then incubated O/N at 4°C with primary antibodies diluted in 0.1% Triton X-100 (Sigma) blocking solution. Slices were stained with the following primary antibodies: Rt anti- CD31 (BD Pharmigen, 1:200), Ms anti-Aβ (BioLegend, clone 6E10, 1:500), Chk anti-GFAP (Aves, 1:2000), Rb anti-cleaved (active) caspase-3 (Cell Signaling, 1:500), Rb anti-CD68 (Abcam, 1:500), Gt anti-IBA1 (Abcam, 1:500), Rt anti-TREM2 (Abcam, 1:200). The following day, the species-appropriate secondary antibodies Alexa Fluor-conjugated (Thermo Fisher, 1:1000) were employed (2hrs at RT), following which 1.5µg/ml DAPI (Invitrogen, D21490) was used as nuclear staining (10’ incubation at RT). Stained sections were imaged with a Nikon Ti2-E fluorescence deconvolution microscope equipped with 340/380, 465/495, 540/580 and 590/650 nm excitation filters, keeping identical settings within each session, and using either 10x or 60x zoom objectives. 60x images were acquired with a 0.5µm Z-stack. In order to have a consistent examination, for each animal, 2 or 3 different images were acquired in the same brain area of interest. To eliminate out of focus signals, all images were deconvolved using the same deconvolution parameters. The analysis was performed in equally thresholded ten–slice maximal intensity projection images, using Fiji, an open source image processing software. Aβ, active caspase-3, GFAP, IBA1, CD68 and TREM2 were measured as positive staining area (number of positive pixels) per acquisition field. Colocalization was analyzed with JaCoP (Just another Colocalization) ImageJ plug-in, calculating Manders’ coefficients (M1 and M2) which imply the actual overlap of the signals (A over B, and B over A, respectively), and represent the true degree of colocalization [145]. M1 and M2 coefficients were scored from 0 to 1 [e.g., M1=1.0 and M2=0.7, in red (signal A)- green (signal B) pair, indicate that 100% of red pixels colocalize with green, and 70% of green pixels colocalize with red]. The M1 and M2 coefficient values were then multiplied by the percentage area of Aβ, active caspase-3, or CD68, accordingly, and plotted.

### Vessel width measurement

Vessels were stained using antibodies against CD31, as described above, in WT and CAI-treated or untreated TgSwDI mice. The average number of visible vessels for quantification in each image of DG and cortex per mouse was 16. To measure the width of the BVs, a line perpendicular to the long axis of the vessel was delineated at its widest point (corresponding to the widest diameter of every visible vessel), and analyzed using NIS Elements Analysis software from Nikon. Histograms show the distribution and frequency of the vessel width (µm) in each group.

### Microhemorrhage staining and quantification

Cerebral slices were blocked for 2hrs at RT in blocking solution (10% NGS-Normal Goat Serum, 1% BSA in PBS), following incubation O/N at 4°C with CD31 primary antibody diluted in blocking solution plus 0.1% Triton-X 1:200. The day after, sections were washed in ice-cold PBS and incubated in anti-Rat 647 secondary antibody (1:1000 in blocking buffer), for 2hrs at RT, to identify vessels. Perls Prussian Blue Staining was then performed on the same slices to detect microhemorrhages, as previously published [146–148]. Briefly, brains were incubated in a solution containing 5% C₆FeK₄N₆ in dH_2_O (Macron) and 10% HCl in dH_2_O, mixed in a ratio 1:1, for 45’ at RT. Following, slices were incubated for 15’ at RT in 0.1% DAB (3,3′ Diaminobenzidine) in 1xPBS (Acros) solution, and then in 0.033% H_2_O_2_ in 0.1% DAB, for another 15’ at RT. DAB enhancement of Perls Prussian Blue staining, as described above, highlights iron accumulation in the cerebral tissue. Samples were mounted and processed in light and fluorescence microscopy. Images were assessed for number of microhemorrhages (DAB+ Prussian blue staining adjacent to a CD31+ blood vessel), in the cortex and DG areas, as well as in meningeal arteries. For each animal, microbleeds were counted in the 10x images in two different sequential slices (in both hemispheres), in 5animals/group, and the numbers were plotted as shown.

### Cerebral blood volume and cerebral blood flow measurement

#### Animals and treatment

CBF and CBV measurements were performed in parallel animal groups at the University of Aarus, Denmark. The experimental procedures were performed according to the regulations of the Danish Ministry of Justice and Animal Protection Committees, with the permit 2017-15-0201-01241. Mice were fed a standard diet, or an ATZ- or MTZ-diet, as described above, following which randomize group scans were performed by researchers blinded to the treatment. As vascular impairment [55] starts early in TgSwDI mice, vascular functionality was assessed in 10/11-month-old mice after a 6-month treatment with CAIs.

#### Surgical preparation and training for awake imaging

For cerebral blood volume (CBV) and cerebral blood flow (CBF) assessment, a chronic cranial window was implanted onto the somatosensory area of the barrel cortex (S1BC), as previously described [149]. Briefly, mice were handled 5 days before surgical preparation to reduce stress during the scanning sessions. All surgical procedures were performed under anesthesia with 1.75/2% isoflurane (induction with 3%) with 100% O_2_ flow. To avoid brain edema, 4.8mg/kg dexamethasone was injected subcutaneously before the surgical procedure. The cranial window of ∼3mm was placed on the cortical area +1.5/2mm antero-posterior and +3mm medio-lateral from bregma. The window was closed with a glass plug, and the edges fixed with cyanoacrylate glue (Loctite Super Glue gel, Loctite®). For restraining the mouse during the imaging session, an in-house metal bar was fixed on the frontal bone using dental acrylic (Meliodent, Germany). Ampicillin (200mg/kg) and carprofen (10mg/kg) were provided for 5 days post-surgery. To avoid stress and improve training sessions, handling continued after surgical preparations. For awake imaging, mice underwent daily training sessions until they reached a total training time of ∼2.5hrs (15’/training session, and for each next session, 15’ are added, until ∼2.5hrs were reached). The training sessions were performed using a replica of the custom-built frame used for optical imaging. The movement was tracked in x-y-z directions with an accelerometer to discard any data with excessive movement. Excessive movement was defined as a signal above or below the mean ± 3 SD in the 3 recorded directions. The accelerometer signal detection was performed using a Powerlab acquisition unit and LabChart 8 software (ADInstruments). The scans for CBF and CBV analysis were performed on the same day and optical table. Before each recording trial, we performed an acclimation test for each mouse to get it used to the stimulation paradigm.

#### Intrinsic optical imaging

Relative change in cortical CBV (rCBV) during functional activation was estimated using intrinsic optical signal imaging (IOSI). This scanning methodology relies on hemoglobin (Hb) absorption [150]. An isosbestic point (Greenlight, 570nm LED) for both oxy- and deoxy-Hb was selected to yield total Hb, considered equivalent to CBV. The window was illuminated with a cold LED light source equipped with a 570 ± 2nm bandpass filter (FB570-10, Thorlabs). Images were collected with a CMOS camera (UI-3280CP Rev. 2, IDS Imaging Development Systems GmbH) at 5 frames per second (fps), with a resolution of 2054 x 2054 pixels (3.45µm/pixel). Subsampling was performed by a factor 2 in both horizontal and vertical dimensions to reduce the file size. Two 20 epochs were performed in each mouse, after the acclimation trial (20 epochs). The methodology for processing and generating the intrinsic signal has been previously reported [151]. Briefly, using a custom-written script in MATLAB (MathWorks), the acquired videos were downsampled to 512 x 512 pixels. First, motion correction of the videos was performed using the SPM12 toolbox, re- aligning all frames to the first frame acquired. Next, the intensity of each pixel was estimated over the entire time course. Finally, the baseline for each pixel was calculated as the mean intensity during the first 5” of each epoch, and whisker stimulation evoked response was calculated as relative intensity change to the baseline.

#### Laser Doppler flowmetry

The whisker stimulation evoked changes in cortical CBF were determined using a Laser Doppler Monitor MOORVMS-LDF (Moor Instruments) for laser-Doppler flowmetry (LDF) through the cranial window. The tip of the LDF probe was positioned ∼0.5mm above the cranial window. 2recordings of 20 epochs were recorded at 10Hz. The signal was processed with a Chebyshev filter using a MATLAB custom-written script. The relative change to the baseline (5”) was calculated for each time point. An average was computed for each trial and considered for statistical analysis.

#### Stimulation paradigm

Functional activation consisted of 10” series of gentle air-puffs (∼1bar) delivered by a custom-built air-puff system. Each puff lasted ∼155ms and was delivered to the contralateral whisker pad at 3Hz.

### Astrocyte area, microglia shape quantification and count

Astrocyte cell area measurement and morphological classification of microglia were performed using Nikon NIS Elements Analysis software. Planes containing GFAP+ and IBA1+ cells were identified by manually scrolling through the Z-stack. Only slices containing identified glial cells with nucleus were processed as a maximum intensity projection to avoid overlapping with cells located in more superficial or deeper layers. Maximum intensity projections of both astrocytes and microglia were thresholded to create a binary mask, and the perimeter was delineated to measure the cell surface area. IBA1+ microglia were counted, and morphologically classified as resting, bushy or amoeboid, based on previously published descriptions [152], using NIS Elements Analysis software from Nikon. Briefly, resting microglia are identified as IBA1+ cells with thin processes and small cell bodies. Amoeboid microglia are characterized by large cell bodies, rounded macrophage-like morphology with no or few processes and are associated with maximal proinflammatory activation, oxidative-free radicals, and microglial apoptosis [61, 62]. They can also actively remove endangered but potentially viable neurons, contributing to brain pathology and neurodegeneration [153, 154]. Bushy microglia present an intermediate activation state and a shape between resting and amoeboid, with intermediate/large cell bodies and thick projections, and is typically correlated with a pro-healing phenotype [155, 156].

### Immunoblotting assessments

For WB assay, cerebral tissue was homogenized in RIPA buffer containing 1X Halt protease inhibitors (Thermo Fisher) (100mg brain/1ml homogenization buffer, 20mM Tris Base, 0.25M sucrose, 5mM EDTA, 1mM EGTA). Homogenates were kept in ice for 30’, and spun at 21000xg for 30’ at 4°C. Supernatants were collected and quantified with BCA method. 20, 50 and 35µg of proteins (for cell lysates, mouse brains and human cortices, respectively) were added with 1x Sample buffer (Thermo Fisher) and 1x Sample Reducing Agent (Thermo Fisher), and boiled (100°C) for 5’. Proteins were fractionated by SDS-PAGE in reducing condition, transferred to 0.45µm nitrocellulose membrane (Bio-Rad), and probed with the following primary antibodies: Rb anti-Carbonic Anhydrase-VB (NovusBio, NBP1-86090, 0.4µg/ml), Rb anti-Carbonic Anhydrase-VA (Invitrogen, PA5-36931, 1:500), Rb anti-Carbonic Anhydrase-II (Invitrogen, PA5- 51598, 0,4µg/ml), Ms anti-APP (Millipore, clone 22C11, MAB348, 1:500), Rb anti-APH-1 (Sigma, PRS4003, 1:250), Rb anti-Nicastrin (Cell Signaling, 3632s, 1:200), Rb anti-ADAM10 (Millipore, AB19026, 1:500), and Rb anti-CD68 (Cell Signaling, 97778, 1:500). As internal loading controls, Rb anti-ATP5a (NovusBio, NBP2-15512, 1:300) was used for mitochondrial proteins, and Ms anti-GAPDH (St Cruz, sc32233, 1:500) was used for non-mitochondrial/total proteins. Membranes were then incubated with the appropriate IRDye secondary antibody (LI-COR, 1:10000) for 1hr at RT, and bands acquired with the Odyssey CLX Infrared Imager (LI-COR). The ratios (protein/GAPDH or mitochondrial protein/ATP5a) were plotted as % of WT (or % of Tg, for hAPP), following quantification (Image Studio Lite Vers 5.2).

### Human subjects

For this study, *post mortem* human brains were provided by the Newcastle Brain Tissue Resource, which is funded- in part- by a grant from the UK Medical Research Council (G0400074), by NIHR Newcastle Biomedical Research Centre awarded to the Newcastle upon Tyne NHS Foundation Trust and Newcastle University, and as part of the Brains for Dementia Research Program jointly funded by Alzheimer’s Research UK and Alzheimer’s Society. Based on pathological *post mortem* examination, the brains were evaluated by the brain bank neuropathologist as CAA, AD with CAA, or healthy (if pathology was absent). We analyzed occipital cortices, the most frequently and severely affected brain areas in CAA, which were flash frozen and stored at −80°C until the biochemical analysis described here.

### hCMEC/D3 cells

Immortalized human cerebral microvascular ECs (hCMEC/D3) were obtained from Babette Weksler (Cornell University) [157]. Cells were grown in endothelial basal medium (EBM-2, Lonza), supplemented with growth factors (Hydrocortisone, hFGF-B, VEGF, R3-IGF-1, ascorbic acid, hEGF, and GA-1000) and 5%FBS, and maintained in a humidified cell culture incubator at 37°C and 5%CO_2._

### Aβ peptides and treatment

Aβ40, Aβ40-Q22 and Aβ42 were synthesized by Peptide 2.0 (Chantilly, VA), as previously described [158]. Peptides were dissolved in hexafluoroisopropanol (HFIP) at a 1mM concentration, incubated O/N to allow the breakdown of secondary structures and obtain monodisperse preparations, [16] and then lyophilized using a Benchtop Freeze Dryer (LABCONCO, Kansas City, MO, USA). Lyophilized peptides were resuspended to a 10mM concentration in DMSO, and dH_2_O was added to achieve a final concentration of 1mM. Prior to the cell culture experimental procedures, peptides were further diluted to the final concentration in 1% FBS EBM-2 for cell treatment.

### Carbonic anhydrase silencing in endothelial cells

Downregulation of CA-VB, -VA or -II was obtained using Ambion® Silencer® siRNA, according to manufacturer’s recommendation. Briefly, hCMEC/D3 were seeded to reach a confluency of 60- 70% by 24hrs. On the day of transfection, cells were treated with Lipofectamine RNAiMAX (Invitrogen) and Ambion® Silencer® siRNA (Life Technologies) at a final concentration of 10µM in Opti-MEM™ Reduced Serum Medium (Gibco, Life Technologies). After 4hrs of transfection, cells were supplemented with complete media for 24hrs. After 24hrs, the transfection media was removed, and the cells were grown in complete medium until the experimental endpoint. The efficiency and specificity of CA-VB, -VA or -II downregulation were tested by quantitative RT-PCR and normalized to Cyclophilin-B expression (Cyp-B). Briefly, 48hrs post-transfection, RNA was extracted using the miRNeasy (Qiagen), and cDNA was obtained using the QuantiTect® Reverse Transcription Kit (Qiagen) according to manufacturer’s instructions.

### qRT-PCR

Quantitative RT-PCR was performed with SYBR™ Green (applied biosystems, Thermo Fisher Scientific) and custom synthesized oligonucleotide primers from Gene Link, using the QuantiStudio 3 system (applied biosystems, Thermo Fisher Scientific). CA-VB: 5’TTCGTTCATCCTTCCGGCAT3’ (F) and 5’TTTTAGGGGGTTGCTTGGCT3’ (R). CA-VA: 5’ACTATCGCCCACTTCAACCC3’ (F) and 5’TCTCTAGGACCTTGTGCCCT3’ (R). CA-II: 5’GAGGGTGAACCCGAAGAACT3’ (F) and 5’GGAAGCTTTGATTTGCCTGT3’ (R). Cyp-B: 5’GATGGCACAGGAGGAAAGAG3’ (F) and 5’AGCCAGGCTGTCTTGACTGT3’ (R). Relative gene expression was calculated utilizing the ΔΔCt method vs. scrambled siRNA (siScr).

### Cell Death ELISA

To assess the extent of apoptosis induced by Aβ in the presence or downregulation of CA-VB, CA-VA and CA-II, fragmented nucleosomes formation was quantified using the Cell Death ELISA^PLUS^ assay (Roche Applied Science), as previously published [38]. Briefly, hCMEC/D3 were transfected with siRNA for 48hrs, at which point the cells were challenged with Aβ42 (10µM), Aβ40 (25µM) or Aβ40-Q22 (25µM), for 24hrs. After the 24hr-treatment with Aβ, the plates were centrifuged for 10’ at 200xg, the cells lysed, and fragmented DNA-histone complexes (nucleosomes, indicating apoptosis) quantified by Cell Death ELISA, according to manufacturer’s instructions.

### Statistical analysis

Prior to statistical analysis, outliers were identified and removed from the original dataset using ROUT method with GraphPad Prism 9.1.0 software. Data were analyzed using ordinary unpaired two-tailed t-test or one-way analysis of variance (ANOVA) test, followed by Tukey’s post hoc test (GraphPad Prism 9.1.0). Differences between groups were considered statistically significant when p≤0.05. In all figures and legends, asterisk (*) and plus (+) symbols denote statistically significant differences. If not differently specified, significant differences versus WT animals are indicated with + symbols, graphs are representative of at least 3 independent experiments and data are represented as means ±SEM. In the legends, the number of experiments *in vitro* or the number of mice per condition (or group) are indicated with “N”, while “n” indicates, for *in vitro* treatments, the number of replicates, and *in vivo,* i) the number of technical replicates (e.g. in WB), ii) the counts (e.g. in MH analysis), or iii) the number of measurements acquired (e.g. in IHC).

Behavioral data were analyzed using two-way ANOVA with group (WT, TgSwDI, Tg+ATZ, Tg+MTZ) and sex (females, males), as between-subject factors. For the analysis of rotarod, body weight was used as covariate. Whenever data were judged to be non-normal by inspection of the quantile-quantile (Q-Q) plot and the Kolmogorov-Smirnov (K-S) test, data were transformed using a transform appropriate to the metric (e.g. log of distance, square root of errors). If transformation did not yield normality, differences between groups were analyzed using the nonparametric Kruskal-Wallis test. Significant main effects or interactions were decomposed using simple main effects and Tukey’s post-hoc comparisons for ANOVA or Dunn’s multiple comparisons for nonparametric tests All statistical analyses were performed using IBM SPSS Statistics v25.

For CBV and CBF, the following parameters were estimated from the time-series: i) maximum response during stimulation (peak), ii) time-to-peak response, iii) area under the curve during the 10” stimulation (A.U.C. Stimulation). We used R version 4.03 to compile the database, to perform statistical analysis and plots. All the statistical analysis of time-series parameters was performed constructing linear mixed models with the package ‘lme4’. To analyze the differences in the estimated parameters, we applied a linear mixed model including group (categorical) as a fixed effect. To account for the correlation in the repeated measurements (2 trials) as well as possible drops out of the data for some mice due to corrupted data, we used an indirect specification of covariance using intercepts for the subject (1|mouse). Gender variability was examined by adjusting each model for gender as a fixed effect. The model to use was chosen using the Maximum likelihood ratio test between the model with the fixed effect gender and the one without the effect in question. Finally, a comparison between groups was performed, changing the reference level of the fixed effect group. ** p<0.01. For rCBV, 11 videos were discarded from analysis due to file corruption (WT = 2, Tg-Ctrl = 2, Tg-ATZ = 2, Tg-MTZ =2). A total of 7 ± 3 epochs of CBV and 7 ± 2 epochs of CBF time series were excluded from analysis due to excessive movement during the data acquisition.

## Author Contributions

SF and EC designed and conceptualized the study. EC, RPR, RVT, BGL, RGH, NLL, FA, LD, EGJ and AM performed experiments and statistical analysis. EC and SF wrote the manuscript. SF acquired funding. MAI, LO, TW and ACM provided essential scientific input and all authors revised the manuscript.

## Acknowledgements

This work was supported by NIH R01NS104127 and R01AG062572 grants, the Edward N. and Della L. Thome Memorial Foundation Awards Program in Alzheimer’s Disease Drug Discovery Research, the Alzheimer’s Association (AARG), the Pennsylvania Department of Health Collaborative Research on Alzheimer’s Disease (PA Cure) Grant, awarded to SF, and by the Karen Toffler Charitable Trust, and the Lemole Center for Integrated Lymphatics research. LD, BGL and AM are supported by P01AG060882 to TW. LD and TW are also supported by P30AG066512 to TW.

## Supplementary Figure Legends

**Supplementary Figure 1:**
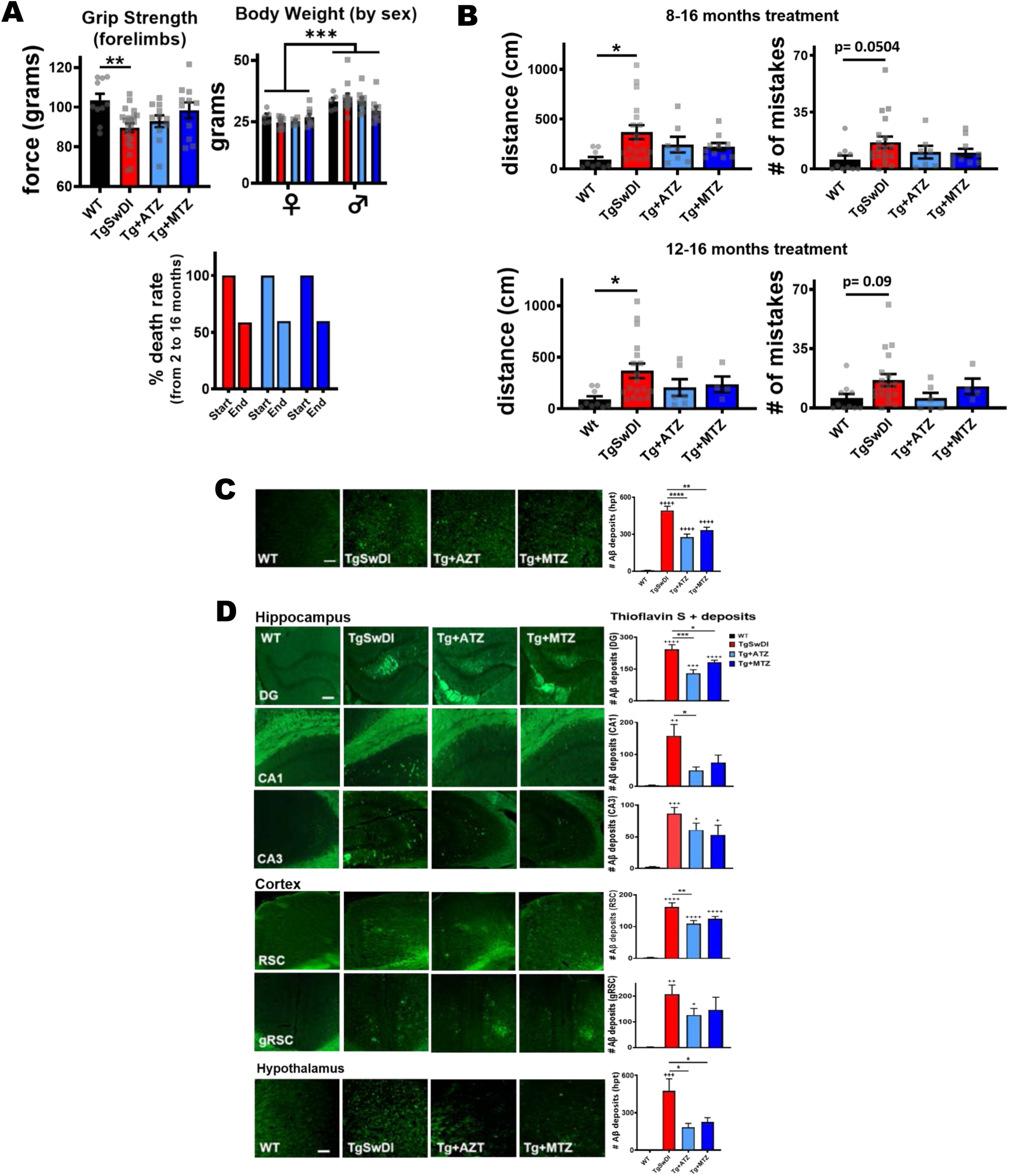
Evaluation of motor function, death rates, spatial memory and Thioflavin S deposits following CAI treatment. **A)** Grip strength of forelimbs (only) showed an impairment in the untreated Tg animals (p=0.009, one-way ANOVA and Tukey). For each group, roughly equivalent numbers of females and males were employed. Females were lighter that males (body weight plotted in grams; p<0.0001), but no significant differences within the same-sex group were observed; moreover, no change in death rates occurred due to ATZ and MTZ treatment, demonstrating that CAIs are not toxic. Animal numbers (after accounting for loss): TgSwDI: N=19, ATZ: N=13 and MTZ: N=14. **B)** Plots show the covered distance (cm) and the number of mistakes to find the escape hole in 16-month-old mice, after 8 months (top graphs; WT and MTZ: N=10, TgSwDI: N=19, ATZ: N=7) or 4 months (bottom graphs; WT: N=10, TgSwDI: N=19, ATZ: N=6 and MTZ N=4). **C)** Representative images of Aβ deposits stained with Thioflavin S in the hypothalamus of 16-month-old mice, treated for 8 months with CAIs. The severe hypothalamic Aβ deposition in TgSwDI was significantly decreased by CAI-treatment. Original magnification, 20x. Scale bar, 150µm. Relative quantification of Thioflavin S+ deposits shown on the right, WT, TgSwDI, ATZ and MTZ N=5, n≥10 measurements acquired/group. **D)** Representative images of Aβ deposits stained with Thioflavin S in the hippocampus, cortex and hypothalamus of 16-month-old mice, treated 4 months with CAIs. Original magnification, 20x. Scale bar, 150µm. Relative quantification of Thioflavin S+ deposits shown on the right, WT, TgSwDI, ATZ and MTZ: N=4, n≥8 measurements acquired/group. * and + p<0.05, ** and ++ p<0.01, *** and +++ p<0.001, and **** and ++++ p<0.0001, One-way Anova and Tukey’s post-hoc test. Data are expressed as mean ± SEM.

**Supplementary Figure 2:**
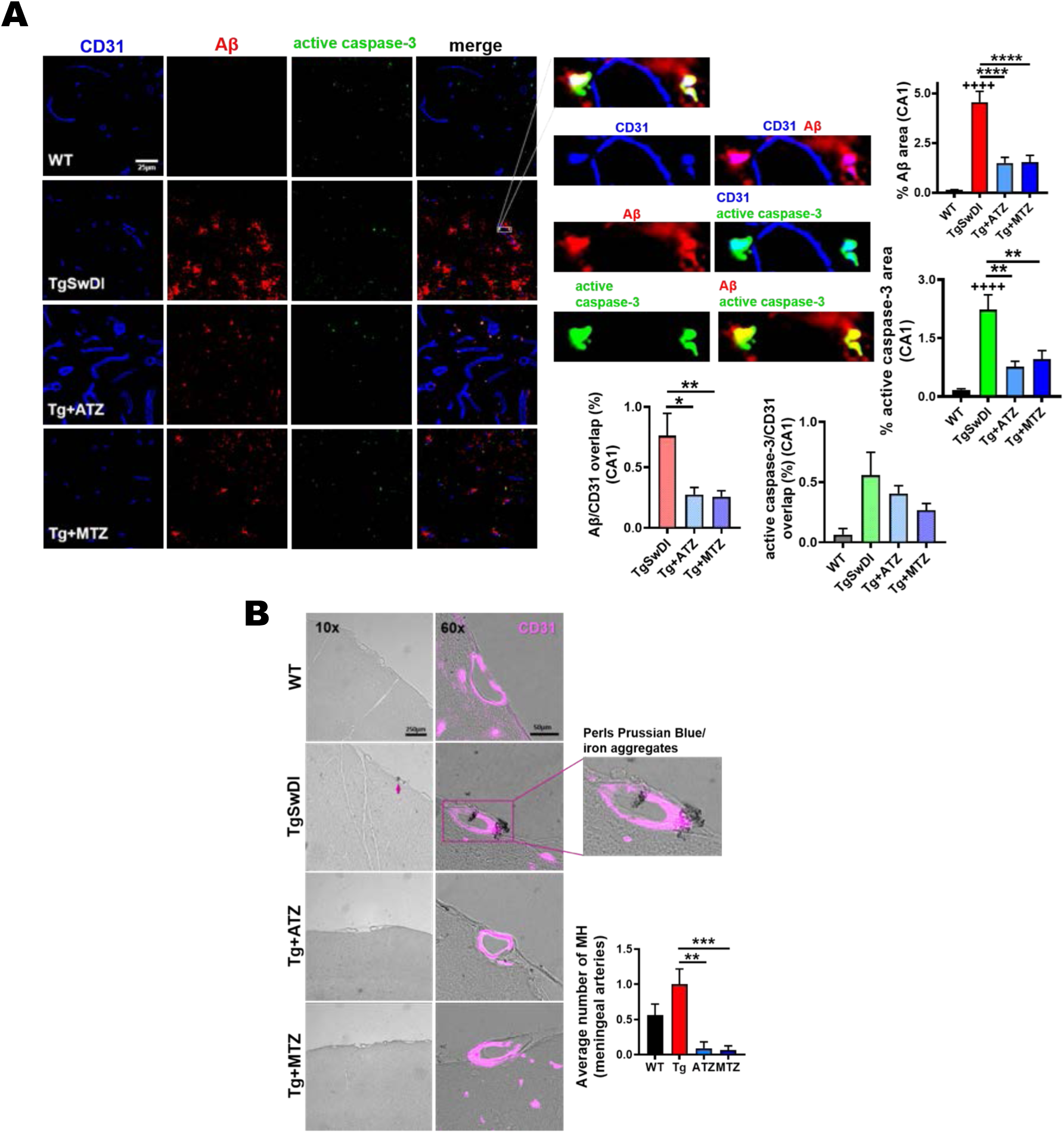
CAIs diminish vascular Aβ burden and endothelial caspase-3 activation in CA1 area in TgSwDI animals. **A)** Representative immunofluorescence images of the CA1 hippocampal region of 16-month-old mice. Compared to WT, untreated TgSwDI animals exhibited massive amount of Aβ (red) and evident caspase-3 activation (green) in CA1, both significantly decreased by 8-month CAI treatment. Original magnification, 60x. Scale bar, 25µm. On the right, the relative quantification is plotted as the percentage of Aβ and active caspase-3 area per acquisition field. For %Aβ, WT, TgSwDI and MTZ: N=5, ATZ: N=4, n≥12 measurements acquired /group. For %active caspase-3, WT, TgSwDI and MTZ: N=5, ATZ: N=3, n≥9 measurements acquired /group. **p<0.01, **** and ++++p<0.0001, One-way ANOVA and Tukey’s post-hoc test. The magnified images show the spatial overlap between the signals: Aβ (red) in the microvasculature; CD31 (blue) (overlap, magenta), endothelial active caspase-3 (green) (overlap, cyan), Aβ overlapping active caspase-3 (yellow). Below, percentage of Aβ and active caspase-3 signals overlapping with CD31 are graphed. ATZ and MTZ significantly attenuate Aβ/CD31 overlap, and show a trend in reducing active caspase-3/CD31 overlap in TgSwDI mice. For Aβ/CD31 overlap, TgSwDI and MTZ: N=5, ATZ: N=4, n≥12 measurements acquired /group. *p<0.05 and **p<0.01, One-way ANOVA and Tukey’s post-hoc test. For active caspase-3/CD31 overlap, =3-5 mice/group, n ≥5 measurements/group. **B)** Representative 10x and 60x images of DAB-enhanced Perls Prussian Blue staining in meningeal arteries of 16-month-old mice, showing higher number of microhemorrhages (MH) (indicated with arrows) in TgSwDI mice, vs WT animals, which are reduced in CAI-treated groups. Colocalization of iron aggregates with CD31+ BVs (magenta) is shown in the 60x and magnified images. Scale bars are 250µm and 50µm, respectively for 10x and 60x magnification. The plot on the right represents the average number of MH counted. WT, TgSwDI and MTZ: N=4, ATZ: N=3, n≥10 counts/group. **p<0.01 and ***p<0.001, One-way ANOVA and Tukey’s post-hoc test. Data are expressed as mean ± SEM.

**Supplementary Figure 3:**
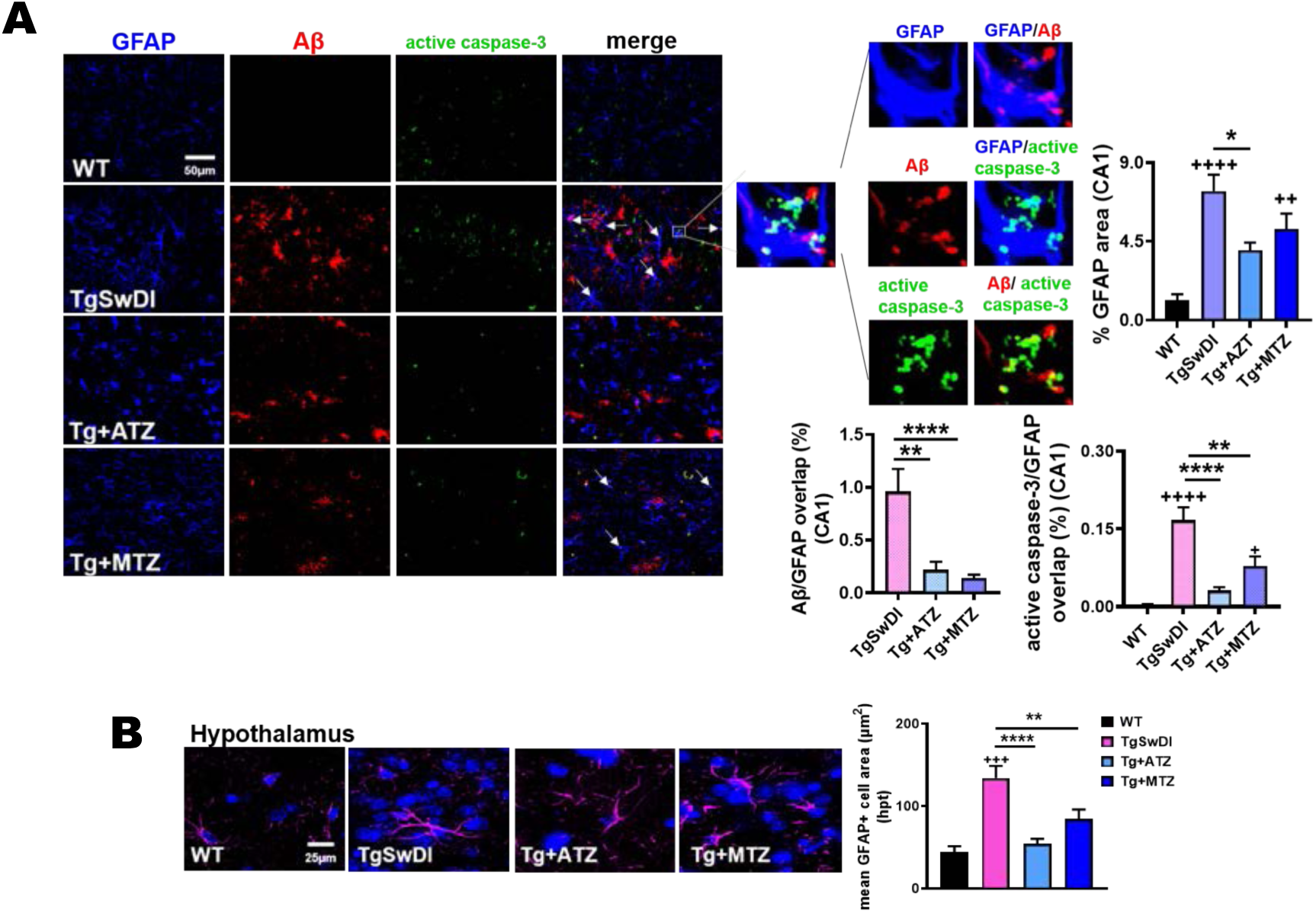
CAIs hamper Aβ deposition and caspase-3 activation in astrocytes, rescuing Aβ-initiated astrogliosis in CA1 area. **A)** Representative immunofluorescence images of CA1 hippocampal area in 16-month-old mice illustrate significant increase of astrocytic marker (GFAP, blue) in TgSwDI animals, compared to WT mice, attenuated by 8-month-CAI-diet. Original magnification, 60x. Scale bar, 50µm. On the right, astrogliosis is plotted as %GFAP area per acquisition field. WT, TgSwDI and MTZ: N=5, ATZ: N=3, n≥9 measurements acquired /group. * p<0.05, ++ p<0.01, ++++ p<0.0001, One-way ANOVA and Tukey’s post-hoc test. The white arrows point to Aβ in astrocytes colocalizing with active caspase-3. The magnified images confirm Aβ (red) content within astrocytes (GFAP) (overlap, magenta), and astrocytic caspase-3 activation (green) (overlap, cyan). Aβ colocalizing with caspase-3, yellow. The graphs below represent the percentage of Aβ and active caspase-3 signals overlapping with GFAP, indicating that both ATZ and MTZ significantly reduce astrocytic Aβ accumulation and caspase-3 activation in astrocytes, in CA1. For Aβ/GFAP overlap, TgSwDI and MTZ: N=5, ATZ: N=3, n≥9 measurements acquired/group. **p<0.01 and **** p<0.0001, One-way ANOVA and Tukey’s post-hoc test. For active caspase-3/GFAP overlap, WT, TgSwDI and MTZ N=5, ATZ N=3, n≥9 measurements acquired/group. + p<0.05, **p<0.01, **** and ++++ p<0.0001, One-way ANOVA and Tukey’s post-hoc test. **B)** Immunofluorescence images represent astrogliosis in the hypothalamus of 16-month-old mice. TgSwDI brains are characterized by increased GFAP+ cell area (µm^2^) (GFAP, magenta), in contrast to age-matched WT, and CAIs attenuate this phenotype. Nuclei stained with DAPI (blue). Original magnification, 60x. Scale bar, 25µm. On the right, the relative quantification, WT, TgSwDI and MTZ: N=5, ATZ: N=4. ** p<0.01, +++ p<0.001, ****p<0.0001, One-way ANOVA and Tukey’s post-hoc test. Data are expressed as mean ± SEM.

**Supplementary Figure 4:**
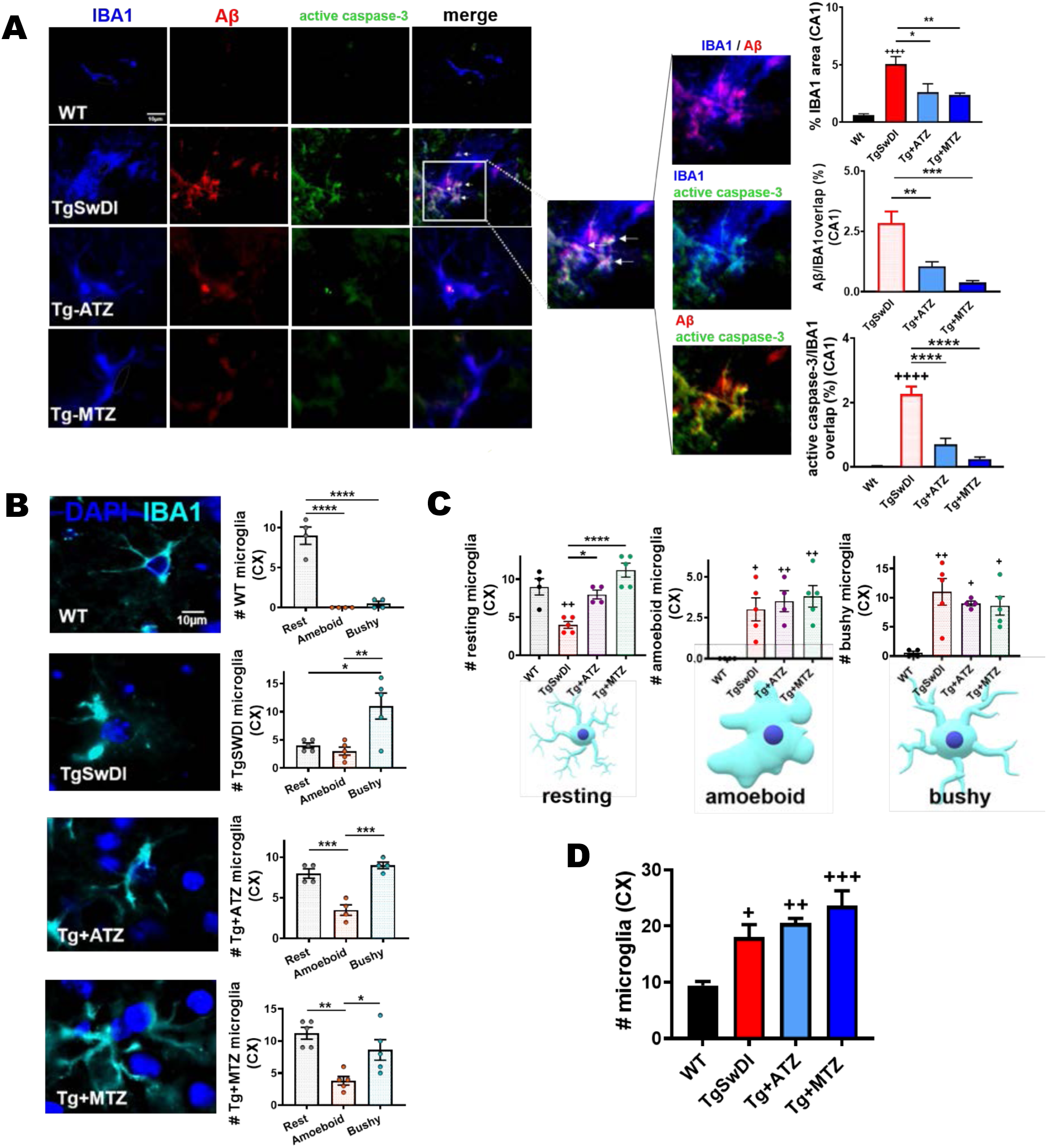
The inhibition of carbonic anhydrases attenuates microglial Aβ accumulation and caspase-3 activation in CA1 area. **A)** Representative immunofluorescence images indicating abundant microgliosis (IBA1, microglia marker in blue), Aβ (red) deposition and active caspase-3 (green) in Tg animals, rescued by CAIs. Original magnification, 60x. Scale bar, 10µm. On the right, microgliosis measured as %IBA1 area per acquisition field. WT, TgSwDI and MTZ: N=5, ATZ: N=4, n≥8 measurements/group. Arrows indicate microglial Aβ accumulation colocalized with active caspase-3. The magnified images show Aβ within IBA1+ microglia (signals overlap in magenta), microglial caspase-3 activity (signals overlap in cyan), or Aβ colocalizing with active caspase-3 (yellow). On the right, plots represent the percentage of Aβ and active caspase-3 signals overlapping with IBA1. For Aβ/IBA1 overlap. WT, TgSwDI and MTZ: N=5, ATZ: N=4, n≥8 measurements/group. For active caspase-3/IBA1 overlap, WT, TgSwDI and MTZ: N=5, ATZ: N=4, n≥8 measurements/group. **B)** Representative immunofluorescence images of microglia (IBA1 marker, cyan) for analysis of resting, amoeboid and bushy morphology. On the right, plots represent the different microglial phenotypes in each treatment group, in cortex. WT mice have resting microglia as the most numerous subpopulation. TgSwDI have bushy microglia as most represented microglial type. ATZ- and MTZ-treated mice have similar amounts of resting and bushy microglia. WT, TgSwDI and MTZ: N=5 ATZ: N=4, n≥8 measurements/group**. C)** Comparison of cortical resting, amoeboid and bushy microglia between the different groups. The number of resting microglia in MTZ- and ATZ-treated mice is significantly higher than in Tg animals, and is similar to WT mice. The amount of cortical amoeboid and bushy microglia does not change in Tg treated and untreated mice. WT, TgSwDI and MTZ N=5, ATZ N=4, n≥8 measurements/group. **D)** Microglia (IBA1+ cells) counts in cortex. TgSwDI and CAIs-treated Tg animals show between 2 and 2.5folds higher number of microglia, compared to WT mice. WT, TgSwDI and MTZ: N=5, ATZ: N=4, n≥8 measurements/group. In **(A-D)**, one-way ANOVA and Tukey’s post-hoc test: * and + p<0.05, ** and ++ p<0.01, *** and +++ p<0.001, **** and ++++ p<0.0001. Data are expressed as mean ± SEM.

**Supplementary Figure 5:**
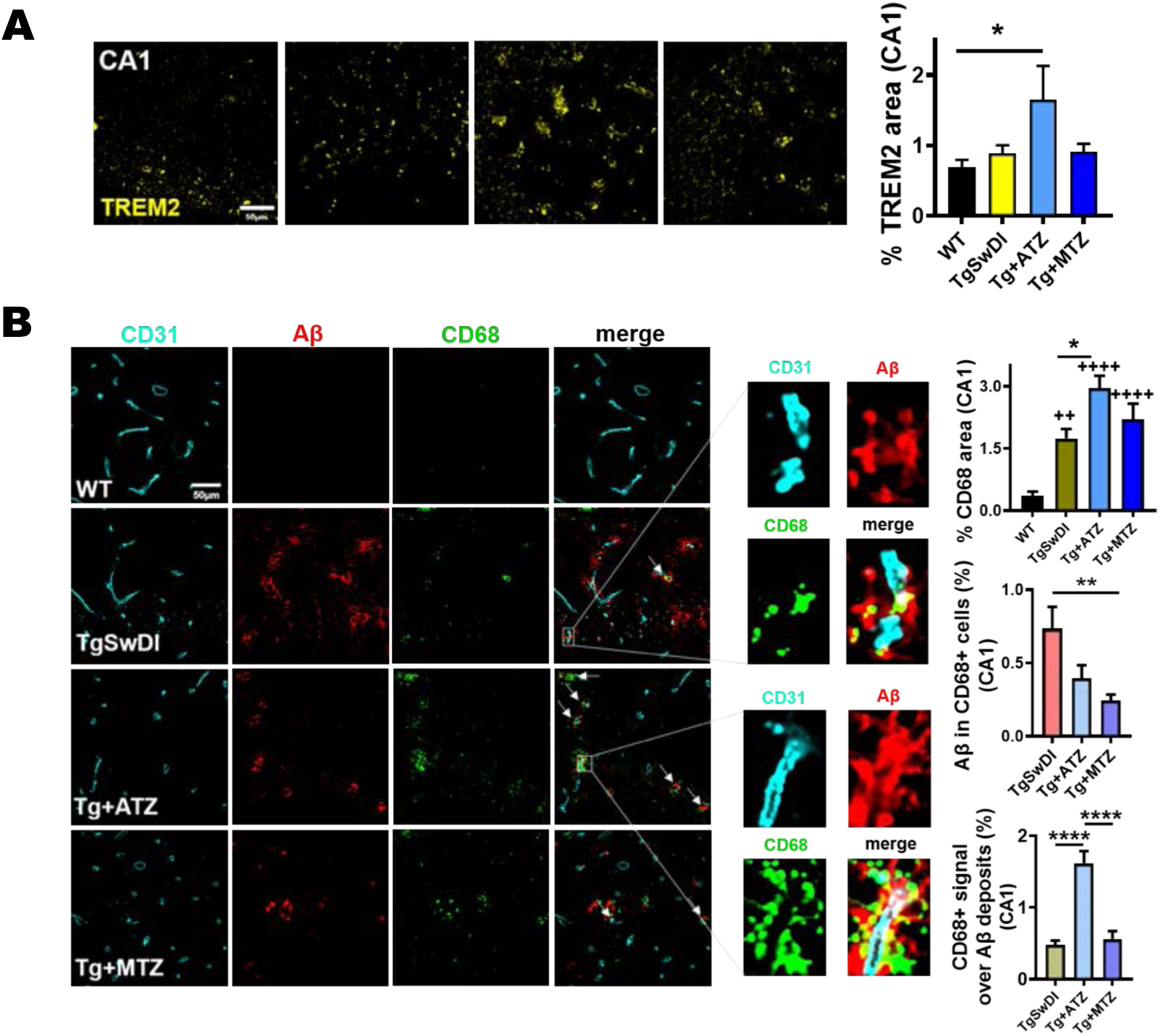
CAIs improve microglial and perivascular macrophage phagocytic activity in CA1 in TgSwDI mice. **A)** Representative IHC images showing that ATZ increases TREM2 expression in microglia in the CA1 hippocampal area. Original magnification, 60x. Scale bar, 50µm. The relative quantification of %TREM2 area is on the right: WT, TgSwDI and MTZ: N=5, ATZ: N=3, n≥9 measurements acquired/group. * p<0.05, One-way ANOVA and Tukey’s post-hoc test. **B)** In CA1 area, CD68 expression (phagocytic activity marker, green) is higher in TgSwDI compared to WT. CAI-treatment boosts CD68 expression, mainly along the cerebral vasculature (stained with CD31, cyan). CD68 is colocalized with perivascular Aβ (red). Original magnification, 60x. Scale bar, 50µm. The plot on the top right shows that CAI-treatment increases perivascular phagocytic activity, measured as %CD68 area per acquisition field. TgSwDI: N=7, WT: N=6, ATZ: N=4, MTZ: N=5, n≥12 measurements acquired/group. * p<0.05, ++ p<0.01, ++++ p<0.0001, One-way ANOVA and Tukey’s post-hoc test. The white arrows indicate Aβ surrounding the microvasculature colocalized with CD68+ perivascular macrophages (PVM), as shown in yellow in the merged magnified images. On the right, colocalization plots for both Aβ within CD68+ cells (Aβ/CD68) and CD68+ cells over Aβ deposits (CD68/Aβ). TgSwDI: N=7, MTZ: N=5, ATZ: N=4, n≥11 measurements acquired/group. For Aβ/CD68 colocalization, ** p<0.01. For CD68/Aβ colocalization, **** p<0.0001, One-way ANOVA and Tukey’s post-hoc test. Data are expressed as mean ± SEM.

**Supplementary Figure 6:**
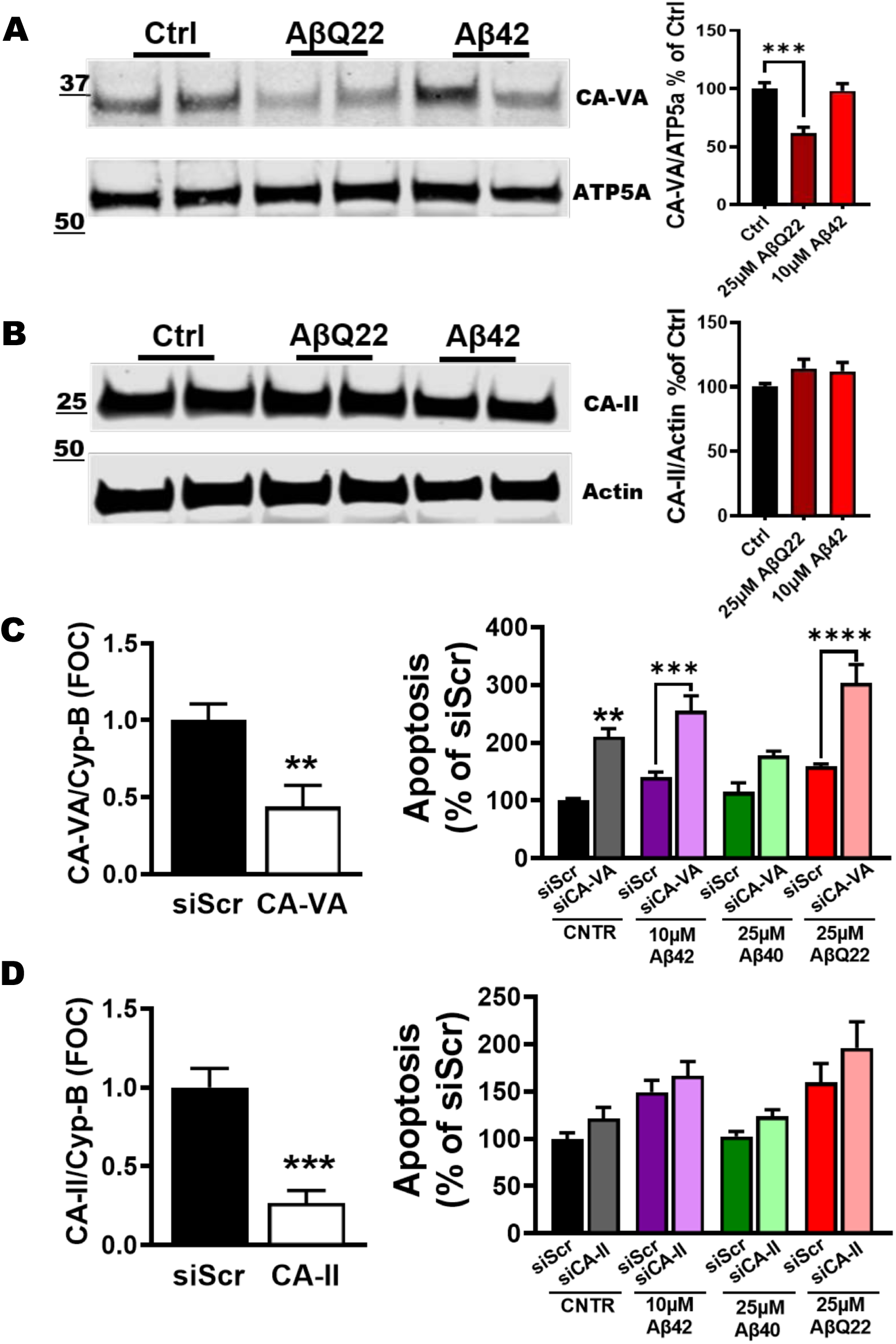
Impact of CA-VA and CA-II downregulation on apoptosis. **A, B)** Expression of CA-VA (**A**) and CA-II (**B**) in hCMEC/D3 cells. Western Blot analysis of CA-VA and CA-II after 24hrs of treatment with Aβ40-Q22 (25µM) and Aβ42 (10µM). CA-VA was normalized to mitochondrial protein ATP5a, while cytosolic CA-II was normalized to actin. Quantification on the right. The expression of CA-VA is significantly reduced following 24hr Aβ40-Q22 treatment. Data represents the combination of at least three experiments, each with 2 replicates, graphed as mean + SEM. Ctrl vs. Q22 ***p<0.001 **C)** qRT-PCR for the mRNA expression levels of CA-VA and Cyp-B (control) at 48hrs post-transfection with siRNA for CA- VA (siCA-VA) or with a scrambled sequence (siScr). On the right, the impact of CA-VA downregulation on apoptosis, as measured by the formation of fragmented nucleosomes, after Aβ42 (10µM), Aβ40 (25µM) or AβQ22 (25µM) challenge for 24hrs. **p<0.01, ***p<0.001, ****p<0.0001 vs. siScr control, One-way ANOVA and Tukey’s post-hoc test. **D)** qRT-PCR for the mRNA expression levels of CA-II and Cyp-B (control) at 48hrs post-transfection with siRNA for CA-II (siCA-II) or with a scrambled sequence (siScr). ***P<0.001 vs siScr, Unpaired two-tailed t-test. On the right, the effects of CA-II downregulation in apoptosis, as measured by the formation of fragmented nucleosomes, after Aβ42 (10µM), Aβ40 (25µM) or Q22 (25µM) challenge for 24hrs. In **C** and **D**, graphs display one representative experiment of at least N=3 experiments, each one performed in duplicate (n=2). Data are expressed as mean ± SEM.

## Notes

### Competing Interest Statement

The authors have declared no competing interest.

### Summary of Updates

Part of the results obtained in the mouse brains have been confirmed in human subjects as well.

